# Divergent organelle allocation in the evolution of sperm gigantism revealed from subcellular quantification of nematode sperm with electron microscopy

**DOI:** 10.1101/2025.05.31.657168

**Authors:** Rebecca Schalkowski, Asher D. Cutter

## Abstract

Sperm gigantism evolved multiple times independently among *Caenorhabditis* nematodes, raising the question of whether intracellular allocation strategies evolved in concert with cell size. Allocation to intracellular components might evolve from direct selection on specific subcellular constituents that affect cell size indirectly, or instead as a byproduct of selection on cell size per se. We used transmission electron microscopy of spermatozoa to quantify investment in pseudopods, mitochondria, and membranous organelles (MOs) from *C. macrosperma* and *C. nouraguensis*, related species divergent in sperm size. We demonstrate that *C. macrosperma* allocates more to mitochondria, in both total and relative terms, consistent with larger sperm cells having greater energetic demands associated with longevity, adhesion, and motility functions. Similar relative pseudopod sizes between species, however, is consistent with an optimal pseudopod:cell body ratio. MO size and distribution patterns within cells implicate *C. macrosperma* having lower relative investment in MO contributions to seminal fluid, thus excluding increased investment in MOs and pseudopod as drivers of sperm gigantism in *C. macrosperma*. We conclude that cell size per se likely represents the primary target of selection in the evolution of sperm gigantism, with mitochondrial traits likely evolving as a consequence of increased energetic demands of giant sperm cells.

## Introduction

Male gametes, sperm cells and pollen, are some of the most diverse cell types in the natural world and the only cells that independently perform their primary function outside their body of origin (Birkhead, Hosken, & Pitnick, 2009). This task must be accomplished in a range of different habitats, e.g., salt water for external fertilizers like sea urchins and fish (Ito et al., 2022; Levitan, 2000; Timourian & Watchmaker, 1970), air for wind pollinated plants (Hall & Walter, 2011; Niklas, 1985), and a broad array of female reproductive tracts that range from benign to antagonistic environments to navigate. To accomplish the goal of fertilizing an egg, some sperm maneuver through anatomically convoluted reproductive anatomies and withstand chemically unwelcoming milieus, as in some insects (Hellriegel & Ward, 1998; Higginson, Miller, Segraves, & Pitnick, 2012; Presgraves, Baker, & Wilkinson, 1999), or evade digestion, as in the female reproductive “stomachs” of some ducks (Harman, 2023). They often compete with sperm of other males in polygamous species but also compete with sperm cells from ejaculates of the same male (Birkhead & Hunter, 1990; Sutter & Immler, 2020), and are subject to female selection among potential fertilizing sperm (cryptic female choice) (Eberhard, 1994; Stumpf, Martinez-Mota, Milich, Righini, & Shattuck, 2011). These diverse challenges that sperm face have resulted in the evolution of spermatozoa across taxa with vast differences in cell morphology and other cell properties.

One common morphological strategy to promote rapid movement toward the egg and past competing sperm cells is increased sperm cell size. In anatomically “typical” sperm made of cell body (head), acrosome, and flagellum (tail), increased size usually refers to a longer tail, as a larger flagellum will increase a sperm cell’s swimming speed and allow it to outcompete other sperm as well as aid in fertilization by penetrating the eggshell and entering through the zona via thrusting motion (Baltz & Cone, 1986). Nonetheless, size-independent alternative evolutionary outcomes across the animal kingdom can involve one or multiple spines and hooks (e.g. *Malacostraca*), multiple or no flagella, and sperm conjugation, among other strategies (Birkhead et al., 2009). In addition to mechanical and morphological diversity, male ejaculates in many species include seminal fluids that accompany the spermatozoa. These seminal fluids often contain peptides and proteins that aid the spermatozoa by (i) providing a habitable environment for the sperm cells, as through altering reproductive tract pH, or by providing a source of energy and immune defence, (ii) outcompeting competitor sperm, and (iii) interacting with female reproductive tissue to stimulate ovulation, facilitate egg fertilization, and “guard” the female by reducing sexual receptivity and/or forming copulatory plugs (Hopkins, Sepil, & Wigby, 2017; Laflamme & Wolfner, 2013; Ramm, 2020; Schjenken & Robertson, 2020; Sirot, Wong, Chapman, & Wolfner, 2015).

An exceptional example of diverse sperm morphology within a single genus is the nematode roundworm *Caenorhabditis*. Sperm gigantism has evolved at least five times independently in this group (Vielle et al., 2016). *Caenorhabditis* sperm cells are amoeboid, in contrast to the “typical” structure of spermatozoa (Justine, 2002; Tarín & Cano, 2012). Their sperm comprise a cell body, containing subcellular organelles and nucleus, and a pseudopod for locomotion (Nelson, Roberts, & Ward, 1982). Like all sperm cells, *Caenorhabditis* sperm are highly derived cells, lacking many of the usual internal components of multicellular eukaryotic cells. During spermiogenesis, the spermatids deposit most cell contents that are not needed for mature sperm function into the residual body (Chu & Shakes, 2013; L’Hernault, 2006). The main remaining subcellular features include the nucleus, mitochondria, membranous organelles (MOs) that are Golgi-derived secretory membranous vesicles, and a large quantity of cytoplasmatic “major sperm protein” (MSP) (Nelson et al., 1982; Nelson & Ward, 1980). MOs release their contents into the seminal fluid after activation during insemination and transfer to the female’s uterus (Kasimatis, Rehaluk, Rowe, & Cutter, 2023; L’Hernault, 2006). Additionally, the MO membrane reshapes and restructures the sperm cell membrane (Achanzar & Ward, 1997; Kasimatis, Moerdyk-Schauwecker, Timmermeyer, & Phillips, 2018; Roberts & Ward, 1982; Xu & Sternberg, 2003). During the activation process of spermiogenesis, sperm cells also develop pseudopod skeletal structures for motility. The pseudopod is densely packed with the cytoskeletal MSP and allows the cell to crawl and adhere to surfaces as well as displace competitor sperm (Hill & L’Hernault, 2001; Nelson et al., 1982; Nelson & Ward, 1980; Thompson et al., 2015). These sperm cells crawl with their pseudopod to the sites of fertilization inside the female, the spermathecae. An increase in sperm cell size in these species translates to a larger cell body rather than an elongated flagellum, meaning the whole cell increases in size, and consequently, increases in crawling speed and ability to displace competitor sperm (Geldziler et al., 2006; Hill & L’Hernault, 2001; C. W. LaMunyon & Ward, 1998, 2002; Craig W. LaMunyon & Ward, 1999; Palopoli et al., 2015; Ting et al., 2014).

One explanation for the repeated evolution of sperm gigantism in *Caenorhabditis* (Vielle et al., 2016) is a functional benefit to sperm size. Sperm size itself could be the primary target of selection, if selection favours speed, adhesion and displacement ability or cellular longevity in these species more than the production of many smaller sperm cells (LaMunyon & Ward, 2002; Palopoli et al., 2015). Evolutionary responses at the cell level might involve disproportionate allocation to pseudopod or to mitochondria, which are essential for energy metabolism via oxidative phosphorylation as well as calcium homeostasis. In some of the smallest sperm cells of the genus, estimates suggest 30- 60 mitochondria per *C. elegans* sperm, which is considered low (Liu et al., 2023; Tsang & Lemire, 2003). Alternatively, sperm gigantism could have evolved indirectly as a consequence of selection favouring a greater quantity of certain subcellular components, with larger cells evolving as a byproduct to accommodate that added volume. For example, MO contents that get released into the seminal fluid upon activation may be most important to the overall quantity of extracellular components to an ejaculate to influence male fertilization success overall rather than through any given sperm cell. A greater investment in MOs, due to collective impacts on seminal fluid, might therefore lead to incidental cell size evolution.

To test whether species with giant sperm invest disproportionately into specific subcellular components, we quantified differences in cell composition between species with giant and standard sized sperm: *C. macrosperma* and *C. nouraguensis*. By comparing intracellular composition of spermatozoa quantitatively from transmission electron microscopy image analysis, we aimed to understand the subcellular causes and corresponding evolutionary pressures responsible for sperm gigantism.

## Methods

### Nematode preparation for sperm cell imaging

We imaged via transmission electron microscopy the activated spermatozoa inside the uterus of mated females for *C. macrosperma* (JU1857) and *C. nouraguensis* (JU1823) as the final developmental form of sperm cells that compete with one another for access to oocytes.

To prepare female nematodes with sperm for imaging, we dissected mated adult females by creating two small incisions on either end of the female, far from the location of the spermatozoa in the uterus to avoid damaging the cells/tissue of interest. Incisions were made using a 25-gauge needle in a watch glass with the chemical fixative 2.5% glutaraldehyde (Electron Microscopy Sciences) in 0.1M Sorenson’s Phosphate buffer (pH 7.4). Samples were then transferred to microcentrifuge tubes and chemically fixed by adding additional 2.5% glutaraldehyde (Electron Microscopy Sciences) in 0.1M Sorenson’s Phosphate buffer (pH 7.4) and incubated at 4°C for 3 days. Following incubation, nematode samples were washed three times with 0.1M Sorenson’s Phosphate buffer (pH 7.4) for 10 minutes each. We then incubated the samples in 1% Osmium tetroxide (OsO4, Electron Microscopy Sciences) in 0.1M Sorenson’s Phosphate buffer (pH 7.4) for one hour, followed by three 0.1M Sorenson’s Phosphate buffer (pH 7.4) washes for 10 minutes respectively. To progressively dehydrate the samples, we washed them three times in an ascending ethanol series in which increasing proportions of 0.1M Sorenson’s Phosphate buffer (pH 7.4) are replaced with EtOH from 50%, 70%, and 90% EtOH for 10 minutes per concentration. To fully replace all water, the samples were washed twice in 100% EtOH for 10 minutes each.

Worms were embedded by progressively displacing the ethanol with Spurr’s resin using decreasing concentrations of EtOH while increasing the concentration of Spurr’s resin as follows. Samples were incubated in 3:1 EtOH to Spurr’s resin for 30 min, before placing them in 1:1 EtOH to Spurr’s resin for another 30 min, followed by one hour incubation in 1:3 EtOH to Spurr’s resin. Lastly, the samples were incubated for one hour in 100% Spurr’s resin and then placed in an oven at 65°C overnight to polymerize and complete the embedding process. On the next day, the samples in resin blocks were removed from the oven and sectioned in preparation for microscopy. We used a Leica EM UC6 ultramicrotome to cut 1 um semithin and 100 nm ultrathin longitudinal sections from the blocks. The semithin sections were heat-fixed and stained with toluidine (1 min) and methylene blue (1 min) to use with light microscopy to determine proximity to target area for ultrathin sections and transmission electron microscope (TEM) imaging. The ultrathin sections were picked up either on 200-mesh grids or collected in Formvar coated slotted copper grids. We stained grids using Uranyless (2 min, Electron Microscopy Sciences) followed by three washes with single drops of distilled water and then Reynolds lead citrate (5 min). After two final washes with drops of distilled water, sections were left to air dry before examination and imaging with a Hitachi HT7700 Transmission Electron Microscope, operated at 80 Ky. For imaging we used an AMT XR-111 digital camera with AMT Capture Engine software. This protocol applied to longitudinal sections of male nematodes, intended to access earlier stages of spermatogenesis, did not reliably achieve high-quality staining and imaging, suggesting that alternative protocols may be superior for this alternate application to our focus on mature spermatozoa in the uteri of mated females. All chemicals were purchased from Sigma Aldrich unless otherwise indicated.

### Image and data analyses

TEM .tif images were analyzed using ImageJ software, after scale bar calibration to measure areas in µm^2^ from regions identified using the freehand selection tool. We measured the cross-sectional area of spermatozoa in terms of the overall cell size (including cell body and pseudopod), the pseudopod, nucleus, individual and combined mitochondrion area per cell, and individual and combined MO area per cell (Figure 1). Additional metrics were estimated from these direct measurements, including peripheral MO density and whole-ejaculate metrics (see below). We tested for differences between the two species using Welch’s t-test, treating each cell as the unit of replication (n*_C. macrosperma_*=22 and n*_C. nouraguensis_*=46). Average counts of sperm cells per ejaculate were inferred from Vielle et al. (2016) and used to estimate MO area differences for sperm cross sections per whole ejaculate. All analyses used the statistical programming language R version 4.2.2 (R Core Team, 2022).

**Figure 1:**
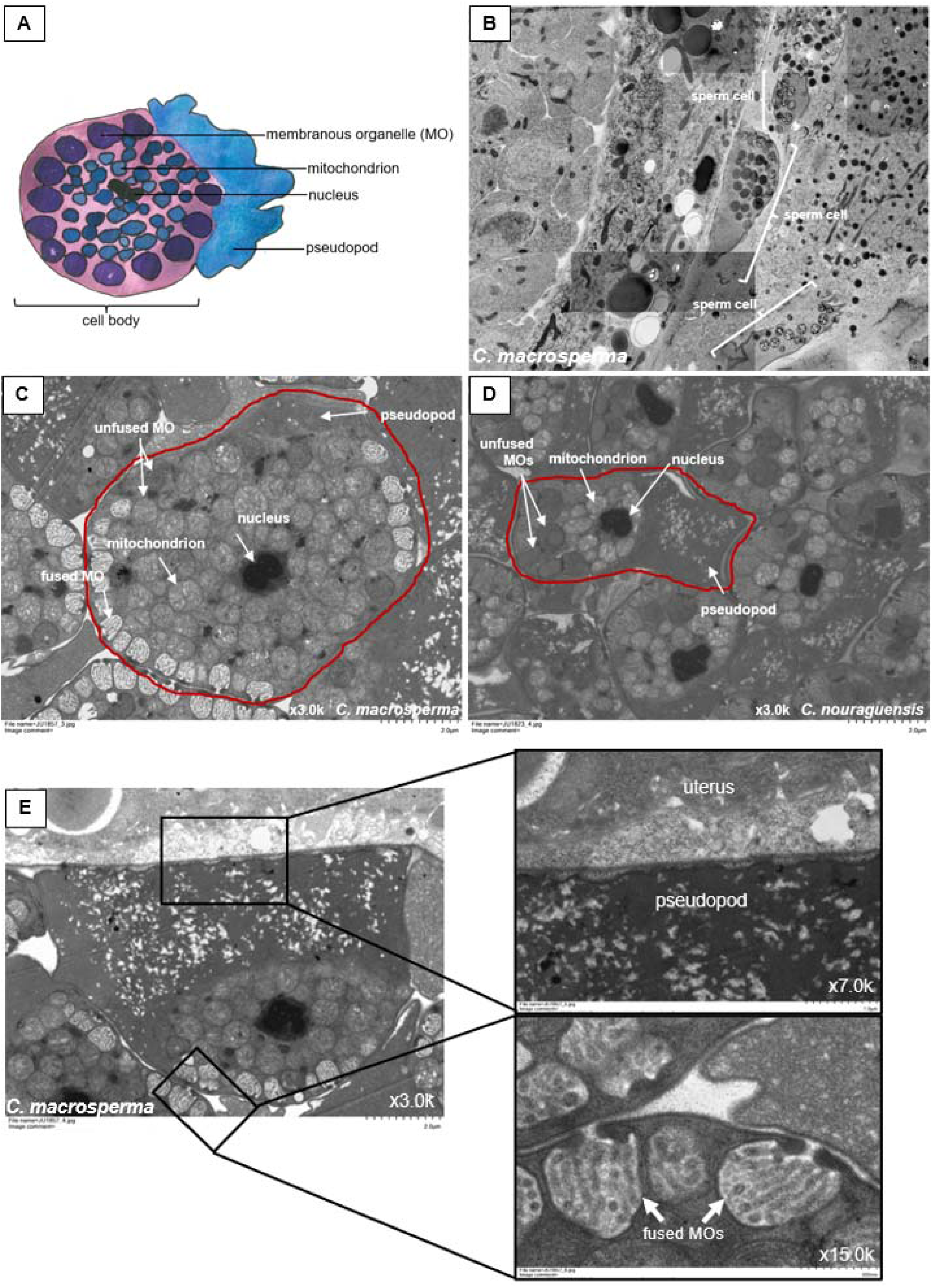
Diagram of activated spermatozoan and cross-sections of mature sperm cells from TEM images. (A) Sperm cell diagram with sperm composition and general location of cell components in an activated sperm cell. (B-E) These images were taken inside a female’s uterus, where sperm is activated. (B) Detailed TEM image of activated *C. macrosperma* spermatozoa inside a conspecific female (courtesy of Ben Mucahy) (C-D) close-up of activated spermatozoa of (C) *C. macrosperma* with spermatozoon in the center surrounded by more (incompletely shown) sperm cells, and at the same magnification (3.0k) in (D) multiple *C. nouraguensis* spermatozoa, illustrating the 4-fold difference in cell size. (E) Close-up of sperm adhesion via pseudopod and fused MOs.

In place of standardizing the MO area in a single sperm cell by the area of that cell, we calculated a metric to better reflect the role of MOs at the sperm cell membrane in restructuring sperm cell membranes and releasing MO contents. Therefore, we estimated MO area relative to sperm cell perimeter from an approximation of cross-sectional length of the cell membrane perimeter by retrieving the circumference of a circle of the same area as the respective sperm cell body area, as cell body perimeter was not recorded directly during measurement. “Peripheral MO area” was estimated as the total MO area (μm^2^) of a cell per unit length (μm) of the circumference of an inferred, idealized circle of the same area as that sperm’s cell body.

We estimated a lower-bound for MO volume per ejaculate from the average number of sperm cells in an ejaculate of a single male for each species (Vielle et al., 2016). We used the mean across cells of a given species to calculate average per-MO size (μm^2^) to estimate the radius of an inferred, idealized circle of the same area to then compute the volume of a corresponding sphere. The product of this hypothetical MO volume, the mean number of MOs in the cross-section of a single sperm cell, and the number of sperm cells in a single ejaculate provided our metric for overall MO volume (*i.e.*, seminal fluid contribution) for each species. The estimated MO volume constitutes a lower-bound because most cells were not thin-sectioned at the widest point of the sperm cell and only the number of MOs in a sperm cell’s 2D cross-section were used in calculations.

## Results

### Cell and pseudopod size of activated spermatozoa in female uteri

Using transmission electron microscopy (TEM), we contrasted *C. macrosperma* giant sperm with *C. nouraguensis* standard-sized sperm for measurements of subcellular components of activated spermatozoa inside the uteri of mated females. First, we confirmed that mature sperm cells from *C. macrosperma* are significantly larger than *C. nouraguensis*, as reported for light microscopy of immature spermatids (Vielle et al., 2016). TEM cross-sectional areas differed between species by ∼two-fold for both cell body alone (P < 10^-06^) and in combination with pseudopod (P < 10^-07^; Figures 2A and 2B). Pseudopod size also was larger on average for *C. macrosperma* (Figure 2C; P < 10^-05^). However, when standardizing pseudopod size by overall cell size, the relative size of the pseudopod did not differ between species in comprising approximately one-third of the spermatozoa cross-sectional area (Figure 2D; P = 0.09 and Figure S1).

**Figure 2:**
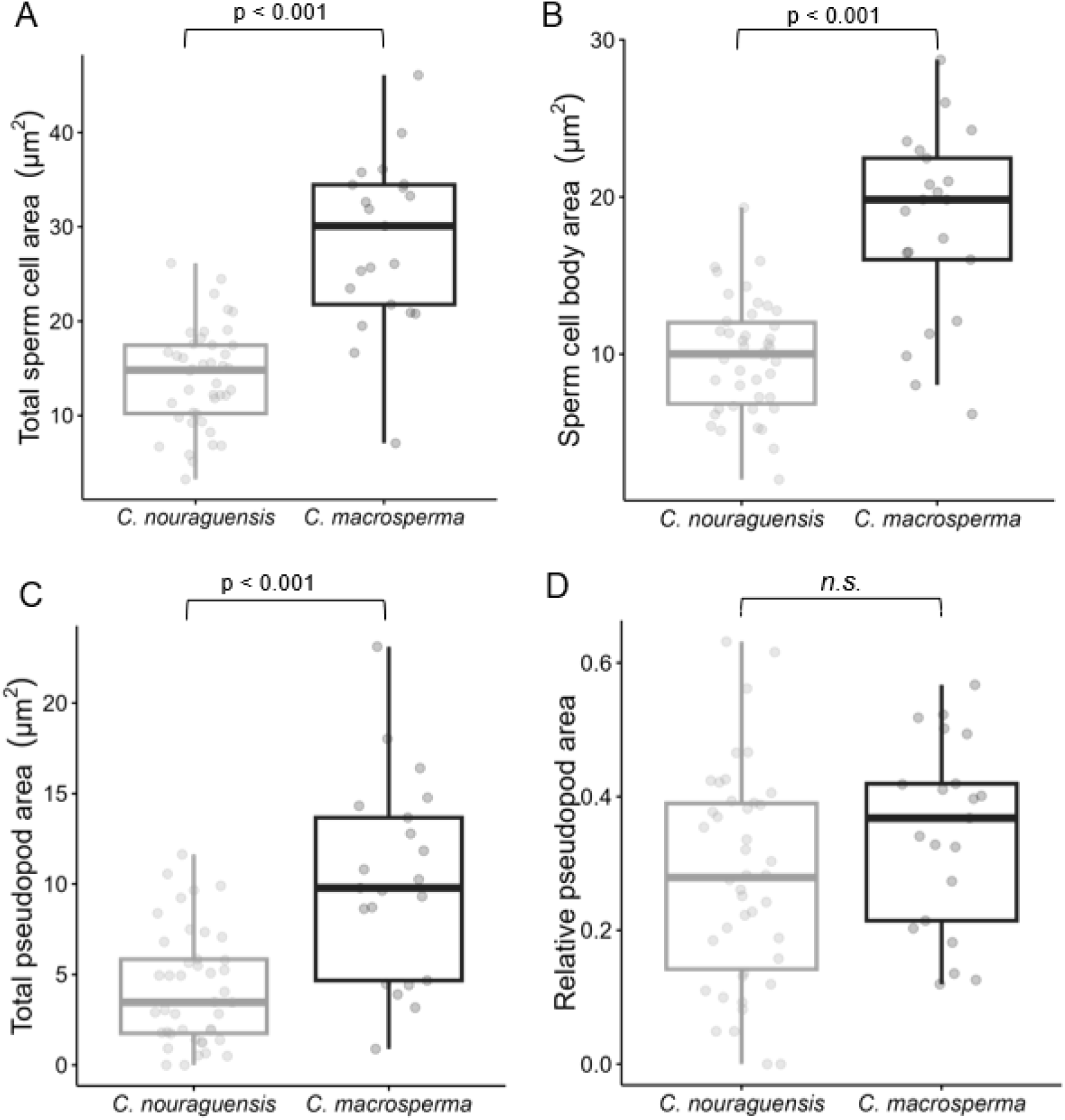
Cross-sectional area of mature sperm cells from TEM images. (A) Area of the total sperm cell including cell body and pseudopod in µm^2^ is greater in *C. macrosperma* (p = 2.7*10^-07^). (B) Sperm cell body area, the sperm cell without pseudopod in µm^2^ is greater in *C. macrosperma* (p = 2.5*10^-06^). (C) Total pseudopod area in µm^2^ is greater in *C. macrosperma* (p = 9.4*10^-05^). (D) Relative pseudopod area standardized per total sperm cell does not differ significantly between species (p = 0.0915). Boxes are superimposed on raw data and show first and third interquartile ranges (IQR), the horizontal bar shows the median, and whiskers range from smallest to largest values within 1.5*IQR respectively.

### Mitochondrial composition within nematode spermatozoa

Looking within cells, *C. macrosperma* sperm each have a greater number of individual mitochondria on average than does *C. nouraguensis* (Figure 3A, P < 10^-06^). The average size of individual mitochondria also is bigger in *C. macrosperma* (Figure 3B, P < 10^-06^). Consequently, sperm in this species show greater total allocation to mitochondria per sperm cell (P < 10^-07^; Figure 3C). When standardized for the overall larger size of sperm cells in *C. macrosperma*, the density of individual mitochondrial organelles per unit cell area does not differ between species (number of mitochondria per cell, P = 0.47), whereas the relative area of a cell allocated mitochondria remains larger in *C. macrosperma* than in *C. nouraguensis* (P < 10^-05^; Figure 3D). This finding holds regardless of whether we standardize for just the organelle containing cell body (Figure 3D), or the total sperm cell including pseudopod (P < 0.01). These observations are consistent with the idea of disproportionate investment in mitochondria due to an increased need for ATP production in larger sperm cells.

**Figure 3:**
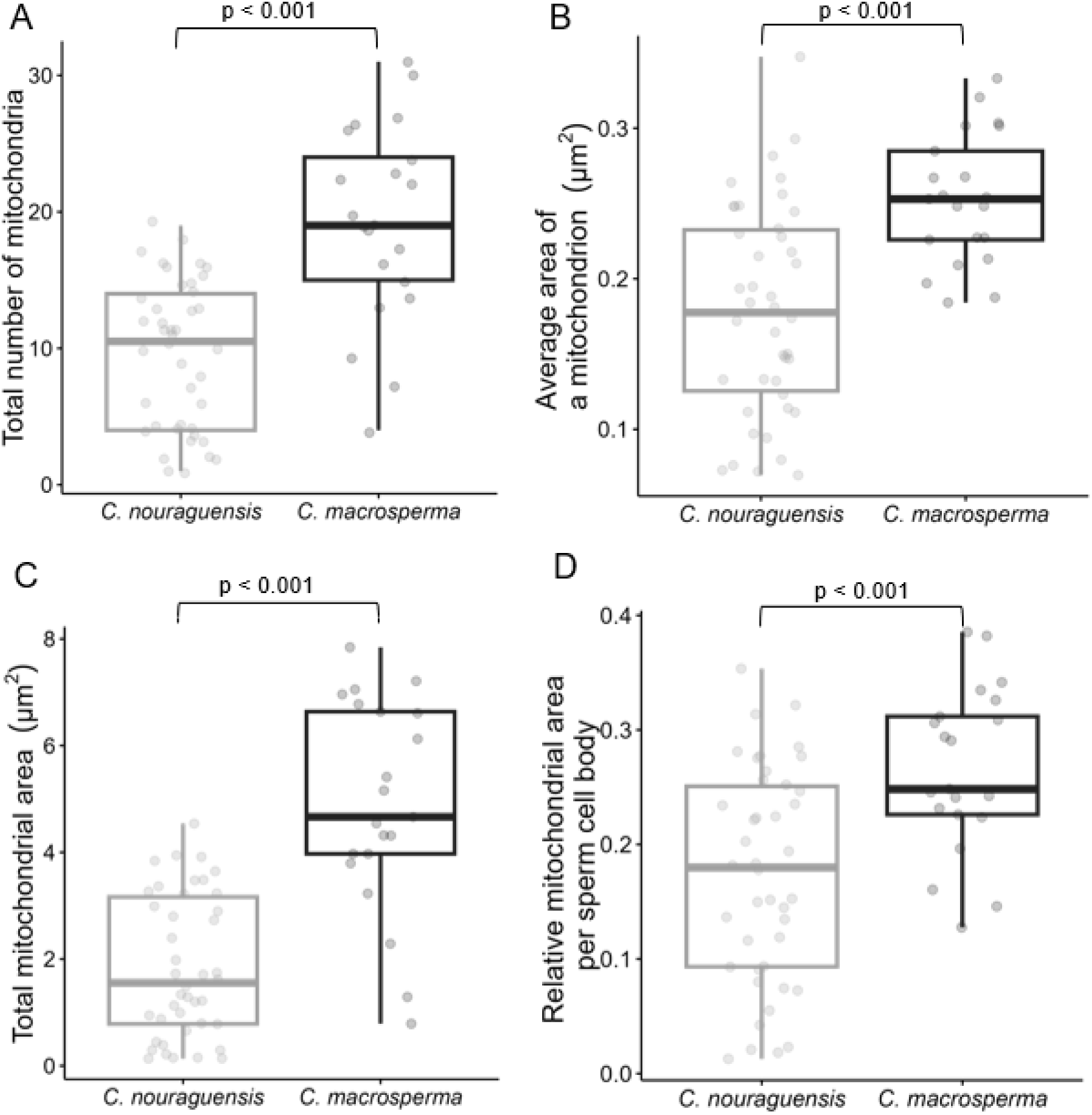
Cross-sectional area of mitochondria from TEM images. (A) The total number of mitochondria is greater in *C. macrosperma* (p = 6.63*10^-06^). (B) The average size of a mitochondrion in µm^2^ is greater in *C. macrosperma* (p = 2.85*10^-06^). (C) Total area of all mitochondria in a cell in µm^2^ is greater in *C. macrosperma* (p = 4.70*10^-07^). (D) Relative mitochondrial area standardized for the size of the sperm cell body is greater in *C. macrosperma* (p = 7.54*10^-05^). Boxes are superimposed on raw data and show first and third interquartile ranges (IQR), the horizontal bar shows the median, and whiskers range from smallest to largest values within 1.5*IQR respectively.

### Membranous organelle (MO) composition in nematode sperm cells and ejaculates

MOs, the organelles that fuse with sperm cell membrane upon activation to secrete their contents into the extracellular seminal fluid, are an important cell trait for sperm competitive function and fertilization success (Achanzar & Ward, 1997). We found the total count of MOs in a given sperm cell to be higher in *C. macrosperma* than *C. nouraguensis* (P < 10^-06^; Figure 4A). However, *C. macrosperma*’s MOs are individually smaller on average than in *C. nouraguensis* (P < 10^-06^; Figure 4B), leading to no difference in the total cross-sectional area allocated to MOs per sperm cell between the species (P = 0.64; Figure 4C). This observation might suggest no difference in the overall investment between species in secretory products contributed by individual sperm.

**Figure 4:**
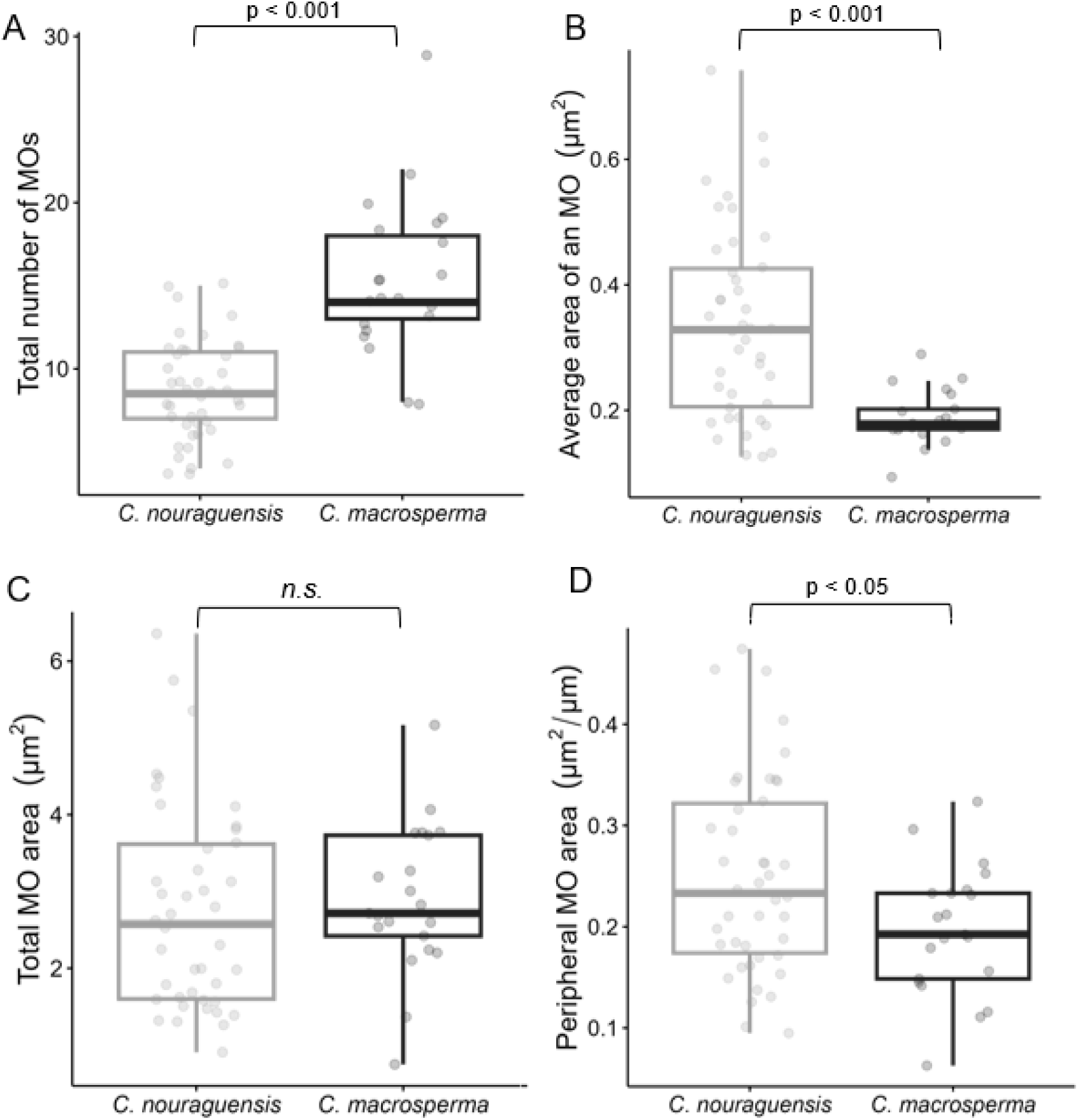
Cross-sectional area of MOs from TEM images. (A) The total number of MOs is greater in *C. macrosperma* (p = 2.22*10^-06^). (B) The average size of an MO in µm^2^ is greater in *C. nouraguensis* (p = 3.08*10^-07^). (C) The total area of MOs in µm^2^ does not differ between the two species (p = 0.642). (D) The peripheral MO area in µm^2^ is greater in *C. nouraguensis* (p = 0.0123). Cell circumference was estimated by taking sperm cell body area and estimating the circumference of a circle of the same area for each cell. Boxes are superimposed on raw data and show first and third interquartile ranges (IQR), the horizontal bar shows the median, and whiskers range from smallest to largest values within 1.5*IQR respectively.

However, when we standardized the cross-sectional area devoted to MOs for sperm cells overall or cell bodies alone, we observed greater relative investment in MOs in *C. nouraguensis* (both P < 10^-05^), the species with smaller sperm. Because MOs need to fuse with the sperm cell membrane to restructure membranes and secrete their contents to contribute to cell performance (Achanzar & Ward, 1997; Roberts & Ward, 1982; Xu & Sternberg, 2003), proximity to and alignment with sperm cell membrane is crucial. To capture this aspect of MO sperm cell functionality, we calculated “peripheral MO area” as MO area per unit length of the perimeter of the cell body (see Methods). We found that peripheral MO area also is greater in *C. nouraguensis* than *C. macrosperma* (P = 0.012; Figure 4D), indicating that MOs occur with higher density along the cell membrane in the species with smaller sperm. The total count of peripheral MOs, however, is greater in *C. macrosperma* than *C. nouraguensis* (P < 0.05), as might result from their smaller individual area in the larger cells of *C. macrosperma*. This implies that the greater density of the peripheral MO component in *C. nouraguensis* sperm cells is achieved by having fewer but larger MOs arranged along a more confined cell perimeter (Figure 1C-E).

The contents of the MO contribute an important component of a male’s seminal fluid (Kasimatis et al., 2018, 2023). The entirety of MO contributions in a male’s seminal fluid will depend on the product of the number of sperm cells and the average MO contribution per cell. We estimate a lower-bound volumetric contribution of MOs to a given ejaculate in *C. macrosperma* of 217.83 µm^3^ to be less than half the volume as in *C. nouraguensis* (603.10 µm^3^). Thus, the relative allocation to MOs per cell and the cumulative allocation of MOs to an ejaculate, presuming that MO size and number contribute proportionately to ejaculation composition, appear greater in the species with smaller and more abundant sperm.

## Discussion

Selection on gamete performance is intense due to the immediate link between gametes and fitness. Diverse adaptations, including sperm cell size, have evolved in response to this evolutionary pressure even among closely related taxa (Gimond et al., 2019; Joly, Korol, & Nevo, 2004; S. Pitnick & Markow, 1994; Scott Pitnick, 1996; Vielle et al., 2016; Birkhead et al., 2009). Theoretical models generally predict that selection should favour numerous small sperm over few larger sperm, given a trade-off between cell size and number (Lehtonen & Kokko, 2011; Parker, Baker, & Smith, 1972). And yet sperm cell gigantism coupled with reduced sperm count has evolved repeatedly in species of *Caenorhabditis* nematodes and other organisms, including *C. macrosperma* (Pitnick & Markow, 1994; Vielle et al., 2016), leaving open questions as to what features contribute to divergence in sperm cell size.

Here, we characterized divergence in subcellular traits of standard and giant spermatozoa in a contrast of two *Caenorhabditis* species to test whether cell size evolution is more consistent with selection on cell size *per se*, with potential subcellular differences in mitochondrial and pseudopod traits, or with indirect selection as might be reflected by the contribution of the quantity of MO contents to the overall ejaculate. Consistent with the direct selection hypothesis, we found that cells from the species with giant sperm (*C. macrosperma*) have more and larger mitochondria resulting in more total and relative allocation to mitochondria, as would be expected from disproportionate investment in cellular energy production. The two species were similar in terms of total MO area and pseudopod size relative to total cell size. However, *C. macrosperma* sperm have smaller MOs that occur with lower density around the cell body periphery, indicative of lower quantitative allocation to the extracellular ejaculate as a whole. Thus, males of *C. macrosperma* evolved to produce fewer but giant sperm cells with disproportionate allocation to mitochondria whereas males of *C. nouraguensis* evolved to produce many small sperm cells with disproportionate allocation to MOs. It remains a challenge to discern which traits are of greatest functional importance in achieving fertilization success under different reproductive conditions.

### Disproportionate allocation to mitochondria with sperm gigantism

The greater total and relative allocation of space to mitochondria in the sperm of *C. macrosperma* could indicate a greater ability of spermatozoa to move fast, to adhere firmly to the uterus wall to secure a spot near the oocyte entrance, and to successfully displace competitor sperm cells, because all these activities require energy from mitochondria-derived ATP. These cell behaviour traits all are characteristics known to increase with sperm size, based on studies of trait variation within species (Geldziler et al., 2006; LaMunyon & Ward, 1998, 2002; LaMunyon & Ward, 1999; Palopoli et al., 2015). Nonetheless, it is unclear how basal metabolic requirements of sperm cells scale with cell size, though metabolic theory as applied from the scale of cells to whole organisms predicts that metabolic rate declines with increasing size (West, Woodruff, & Brown, 2002). Among cells from the same tissue type, larger cells have reduced metabolic activity (Savage et al., 2007). Moreover, there are fundamental differences in allometric scaling between eukaryotes and prokaryotes and between single- and multicellular organisms (DeLong, Okie, Moses, Sibly, & Brown, 2010). *Caenorhabditis* are multicellular organisms, but sperm cells perform their function detached from any somatic tissue and so might be more comparable to single celled organisms, such as yeast or other eukaryotic protists. In yeast, mitochondria increase isometrically (linearly) with cell size (Rafelski et al., 2012), but how that translates to changes in organelle function or cell metabolism is still unknown (Miettinen & Björklund, 2017). Future research will benefit from testing ideas from metabolic theory with sperm cells as well as assessing other features of their biology, for example, whether disproportionate increases in mitochondrial activity with cell size for *Caenorhabditis* spermatozoa might mitigate the otherwise expected allometric decline in metabolism or provide energy to extend sperm cell lifespan in the female reproductive tract.

Even in species where larger sperm size is inherently advantageous in sperm competition, there may be limits on the evolution of size imposed by mitochondrial activity. While most organelle and protein contents increase linearly with cell size, mitochondrial activity is maximized at an intermediate cell size in most multicellular organisms (Miettinen & Björklund, 2016, 2017). Metabolic rate is almost entirely dependent on mitochondria, and metabolic rate also determines cell growth rate (von Bertalanffy, 1934), making mitochondria a major mechanism for cell size determination by growth (Miettinen & Björklund, 2017). However, upper limits to cell size are influenced by diffusion times of nutrients and oxygen in progressively larger cells with greater volume to surface area ratios that limit biochemical reactions (Soh, Banaszak, Kandere-Grzybowska, & Grzybowski, 2013). Often, extraordinarily large cells have elongated morphologies (e.g., neurons) that allow mitochondria to remain close to the cell membrane for gas and nutrient exchange. Function constrains the amoeboid cell morphology of *Caenorhabditis* spermatozoa, however, with a spheroid cell body around the space closest to the cell membrane monopolized by MOs, leaving mitochondria in the center of the cell lumen away from cell membrane. Perhaps this organellar distribution could impose a limiting factor in the evolution of even greater sperm gigantism.

Alternatively, greater connectivity of mitochondria, via mitochondrial fission and fusion, could enable particle transport for metabolism faster than diffusion from the rapid movement of nutrients and oxygen along organelle membranes (Aon, Cortassa, & O’Rourke, 2004; Glancy et al., 2015). Highly connected networks of mitochondria can transport energy from areas close to the cell surface to areas where energy is needed (Miettinen & Björklund, 2017), such as the pseudopod of spermatozoa, which is devoid of mitochondria. Through increased mitochondrial connectivity, mitochondrial dynamics may offset energy transport limitations to allow for larger cells (Miettinen & Björklund, 2016). Recently, mitochondrial exocytosis from *C. elegans* sperm cells was described, with such “mitopherogenesis” proposed as a novel mechanism for secretion of excess mitochondria by sperm cells in the regulation of mitochondrial abundance (Liu et al., 2023), indicating the potential for dynamic mitochondrial regulation within *Caenorhabditis* sperm. Sperm cells are especially small in *C. elegans*, and comparable in size to those of *C. nouraguensis* (LaMunyon & Ward, 1998; Vielle et al., 2016). While *C. macrosperma*’s giant sperm cells have more and larger mitochondria, it remains to be determined whether *Caenorhabditis* species differ in mitochondrial connectivity within or secretion from their sperm cells in ways that could facilitate or constrain sperm cell size evolution.

### Reduced relative allocation to MOs with sperm gigantism

Fusion of MOs to the sperm cell membrane is necessary and required for nematode male fertility, with no known purpose of the minority of unfused MOs within sperm cells (Achanzar & Ward, 1997). The secreted contents of MOs that contribute to seminal fluid, as well as the additional MO-membrane that fuses with the sperm cell membrane, provide this organelle’s main contributions to sperm fitness (Achanzar & Ward, 1997; Kasimatis et al., 2018; Roberts & Ward, 1982; Smith & Stanfield, 2011, 2012; Xu & Sternberg, 2003). Because the surface area of a sphere, or sperm cell membrane, does not increase as rapidly as the volume it contains, however, ever more allocation of MOs to sperm will not fit along the sperm cell body’s membrane.

We predicted that MO abundance would scale with either sperm cell size or total ejaculate size, to the extent that the amount of extracellular secretion by MOs contributes volumetrically-important seminal fluid by an inseminating male to enhance fertilization success. While *C. macrosperma*’s giant sperm cells have more MOs in absolute abundance, they are individually smaller to yield the same relative allocation to MO space within a cell and a lower areal density at the cell periphery than in *C. nouraguensis*. *C. macrosperma* also transfers substantially fewer spermatozoa per ejaculate than *C. nouraguensis*, on average (Vielle et al 2016). It is possible that the fused portion of the larger number of MOs in *C. macrosperma* together contribute disproportionately to the total surface area of the sperm cell plasma membrane, compared to *C. nouraguensis*, though this hypothesis would require further testing. In our extrapolation of MO volume from TEM sections, we estimated that a given male ejaculate from *C. macrosperma* has less than half the combined MO volume of *C. nouraguensis*, and thus less than half the extracellular MO contribution to seminal fluid. Such different investments in MOs at the level of cell and ejaculate might influence sperm competition strategies and outcomes due to how cellular and extracellular components link to sperm function (Achanzar & Ward, 1997; L’Hernault, 2006). However, it is important to note that the qualitative protein composition and activity of secreted MO contents may not scale directly with volume.

Despite the importance of MO fusion to the cell membrane for male fitness, unfused MOs get transferred to the oocyte during zygote formation and, unlike most other sperm cell contents, only get degraded later during early embryonic cell divisions (Al Rawi et al., 2012). It is thus conceivable that MOs and their contents contribute functionally to the developing early embryo or that degradation products provide raw materials for embryo development. This possibility could contribute to speculative explanations for the evolution of sperm gigantism, such that giant sperm cells arose as an evolutionary result of interparental conflict or paternal care, perhaps in the form of energetic or nutritional contribution to zygotes as an unusual kind of a nuptial gift that contributes nutrients via transferred sperm-derived organelles (Al Rawi et al., 2012; Kanzaki et al., 2018; Sato & Sato, 2012; Vielle et al., 2016). Given that only about 70% of MOs fuse to the cell membrane in wild type *C. elegans* (Ward, Argon, & Nelson, 1981), it will be valuable to explore MO fusion more explicitly in species like *C. macrosperma* that have evolved sperm gigantism, and to estimate the quantity of MO membrane contributed to sperm cell membrane (Roberts & Ward, 1982; Xu & Sternberg, 2003). Future work that tracks MO components in embryos could address potential paternal contributions to the embryo, e.g., using techniques that employ heavy carbon labeling of males (Findlay & Swanson, 2010; Findlay, Yi, MacCoss, & Swanson, 2008) or single cell RNA sequencing could test for gene expression products that might get delivered to zygotes (Hotzy et al., 2022; Jimenez-Gonzalez et al., 2023), especially in conjunction with single-molecule fluorescence in situ hybridization (FISH).

Although we have highlighted possible adaptive explanations for the evolution of sperm gigantism, in principle, non-adaptive processes also could contribute. For example, greater relative investment in MOs might be unnecessary for male fitness in *C. macrosperma* if they experience smaller mating group sizes and lower male-male competition that reduces selection on ejaculate components contributed by MOs (McDonald, 2023). Relaxed selection on fertilization might lead to sperm cell size evolution that departs from “optimal” allocation of sperm size and number found in most other *Caenorhabditis*. Mathematical models show how accelerated DNA sequence evolution also can result from sex-limited gene expression due to a weaker strength of selection rather than sexual adaptation (Dapper & Wade, 2016, 2020).

### Allometric pseudopod proportions across sperm cell sizes

The pseudopod, while larger in *C. macrosperma*, appears to scale isometrically with total cell size, suggesting that the ratio of cell body to pseudopod is consistent between species with different average sperm cell sizes for optimal function. This finding is consistent with previous reports (Vielle et al., 2016). The primary functions of the pseudopod are cell motility and adhesion, allowing the sperm cell to move amoeba-like toward the oocyte inside the female reproductive tract, displace competitor sperm, and secure access to those oocytes entering the spermatheca by positioning close to the most distal point of the reproductive tract (L’Hernault, 2006; Smith, 2006). Pseudopod motility depends on major sperm protein (MSP), not actin-myosin, which creates treadmilling forces through dynamic dimerization, breaking down, and re-dimerizing (Miller et al., 2001; Nelson et al., 1982; Smith, 2006). MSP abundance is rate-limiting and the size of the pseudopod depends on the quantity of MSP that is packaged into the cell during spermatogenesis (Bottino, Mogilner, Roberts, Stewart, & Oster, 2002; Burke & Ward, 1983), consistent with demonstrations that overall sperm cell size gets defined early in spermatogenesis (Bottino et al., 2002; Burke & Ward, 1983; Gimond et al., 2019; Vielle et al., 2016). The early developmental determination of both total cell size and pseudopod size suggests that pseudopod to cell ratios are conserved for optimal cell performance. Should other taxa reveal different pseudopod to cell ratios, it may reflect different selective pressures on sperm cell adhesion versus motility, and perhaps may impose different selection for energy metabolism in the allocation of mitochondria within the cell.

### Conclusions

The comparative analysis of subcellular sperm composition using cross-sectional EM imaging in this study leads to the conclusion that selection on sperm size *per se*, rather than indirect selection on organelle features, is most likely responsible for the evolution of sperm cell size differences among *Caenorhabditis* species. The disproportionate allocation of cell size to mitochondria, rather than pseudopod or MOs, in the species with sperm cell gigantism implicates energy metabolism as a key feature in the evolution of sperm cell size in these animals. A powerful extension of the present analysis will be to characterize all *Caenorhabditis* species with sperm gigantism to perform replicated phylogenetic contrasts, and to link metabolic theory directly to sperm cell size variation. In particular, application of 3D EM reconstruction of complete sperm cells, as being conducted for neurons in *C. elegans* (Mulcahy et al., 2018; Witvliet et al., 2021), may more accurately assess volumetric subcellular allocations and reliably quantify fused and unfused MOs to generate deeper insights into the mechanisms and evolution of sperm gigantism.

## Supporting information

data and analysis scripts

## Data availability

Sperm cell and sperm cell component area data for *C. macrosperma* and *C. nouraguensis* is available in Appendix S1. Raw data and R script are published on Zenodo: https://doi.org/10.5281/zenodo.15556660. Nematode strains are available from the authors upon request or from the Félix Lab Strain Database via https://justbio.com/tools/worms/search.php.

## Author contribution

Rebecca Schalkowski and Asher D. Cutter devised the project. RS collected and analyzed the data. RS wrote the manuscript with the help of ADC.

## Acknowledgements

We thank Audrey Chong of the Imaging Facility in the Department of Cell and Systems Biology at the University of Toronto for her technical expertise in the sample preparation and the use of the Hitachi HT7700 Transmission Electron Microscope. We further thank Clotilde Gimond her help with preparing and trouble-shooting nematode dissection techniques, Christine Rehaluk for helping with data collection and image processing, and Katja Kasimatis for feedback on a draft of the manuscript. ADC is supported by Discovery Grant funds from the Natural Sciences and Engineering Research Council of Canada.

## Supplement

### Supplementary Figure

**Supplementary Figure 1:**
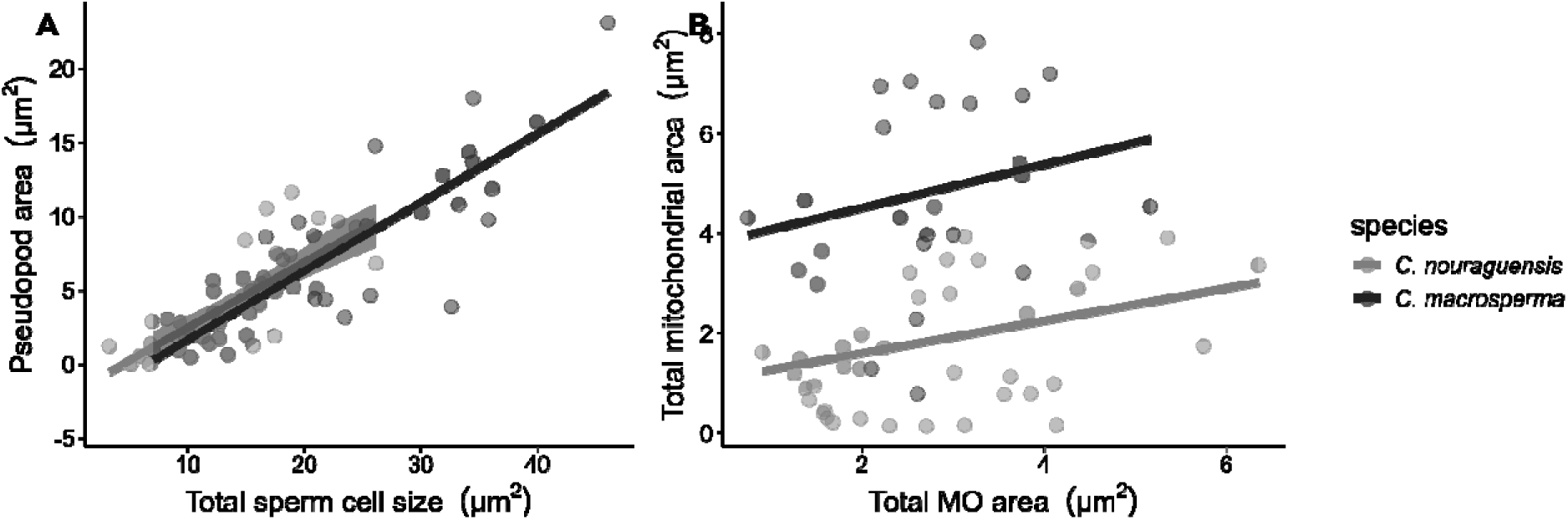
Biplots. **(A)** Pseudopod size relative to total sperm size in µm^2^ from cross-sectional TEM images of spermatozoa. **(B)** Total organelle size showing mitochondria against MO area in µm^2^ from cross-sectional TEM images of spermatozoa. Solid lines represent fitted regression lines and shaded areas indicate 95% confidence intervals for the mean predicted values.

### Formvar-coating of slotted grids for TEM sample preparation

A clean microscopy slide was carefully inserted into a 1% formvar in chloroform solution using a dipping/casting contraption. After drying for a few seconds, the edge of the formvar coat was carefully scored with a razor blade on all sides allowing the formvar film to lift and float on the surface tension of the water when the slide was carefully inserted into the water bath. The submersion in water should occur at a shallow 45-degree angle and the film could be observed to separate from the microscopy slide. From the surface of the water the formvar film was picked up using a dry metal rack with holes slightly larger than the grids. After placing the slotted grids on a hole covered with formvar film, the whole rack was left to dry overnight before careful separation of the slotted grids. We used these grids for most samples after initial trials with mesh grids caused difficulties seeing whole sections in the TEM.

### Test statistics

Statistical tests were Welch’s t-tests comparing *C. macrosperma* and *C. nouraguensis*. Below are the results of all statistical analyses done in the exploration of this data set. No correction for multiple testing was performed due to the exploratory nature of the analysis.

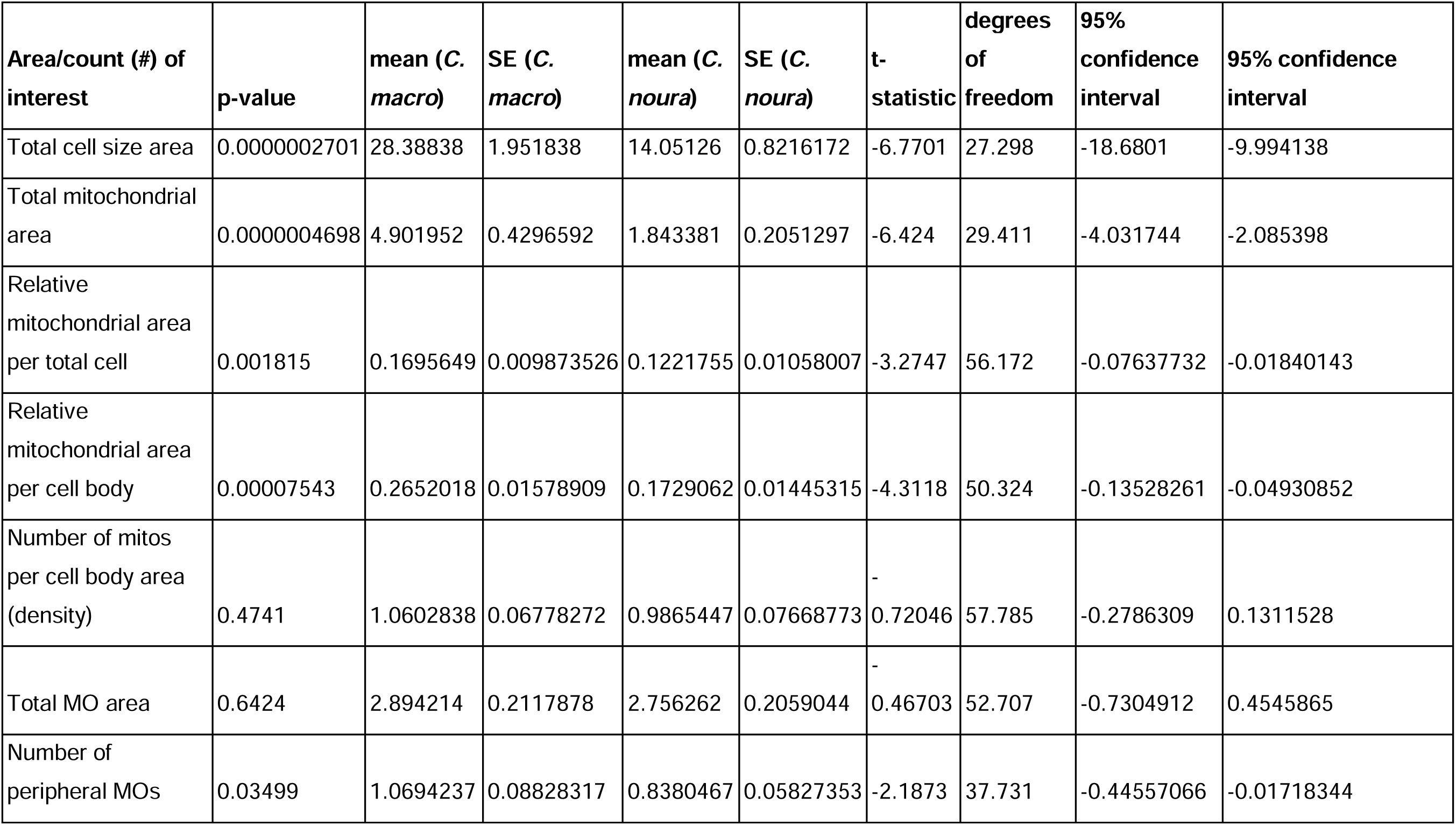
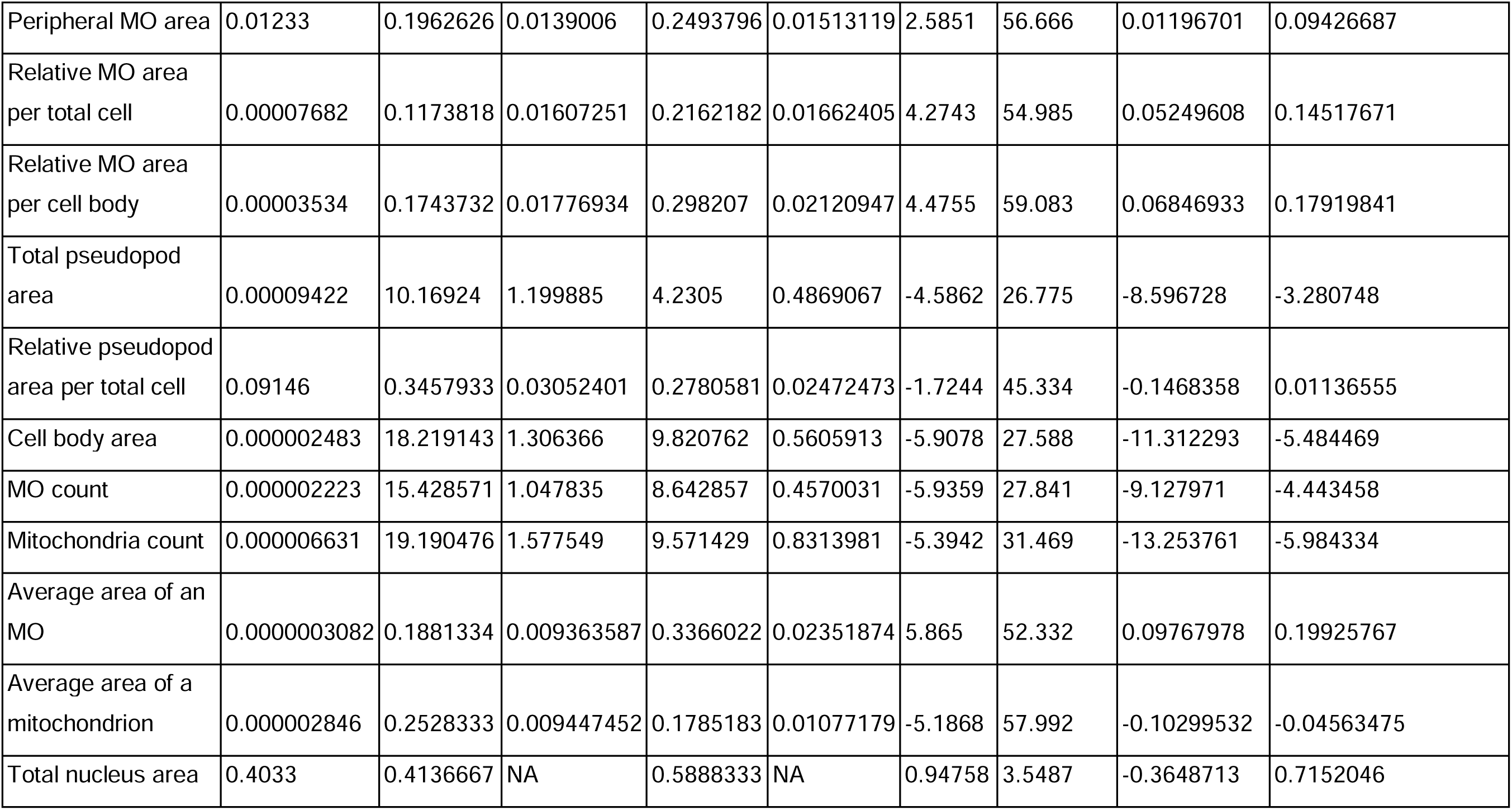

