## Supplementary material for "Divergent organelle allocation in the evolution of sperm gigantism revealed from subcellular quantification of nematode sperm with electron microscopy": data and analysis scripts: README.txt

TEM sperm cell and organelle area analysis in C. nouraguensis and C. macrosperma:

The purpose of this study is to explore differences between sperm cell composition in species with giant vs. standard sized sperm. 
To accomplish this, mated females with sperm in their uteri of the species C. nouraguensis and C. macrosperma were chemically fixed, embedded, thin-sectioned, stained, and TEM imaged. 
This allows us to view cross-sections of sperm cells inside the uterus and, using ImageJ software, measure the area of the total sperm cell, pseudopod, and organelles such as mitochondria, membranous organelles (MOs), and in some cases nuclei. 
Additionally, we can compare relative area of a sperm cell allocated to an organelle type as a proxy for certain investment into that organelle and its function in the sperm cell.

Associated files:
1 - SpermCellAreaDataRecordedValuesOnly.csv
2 - TEMms.Rmd

1 - SpermCellAreaDataRecordedValuesOnly.csv column names & contents:

species - nematode species (C. nouraguensis or C. macrosperma)
specieslabel - refers to label on physical microscopy slide identifying the species and specific sample that was imaged 
unique_ID - column with unique ID that combines all info required to track a specific sperm cell within a specific worm
grid_id - label corresponding to the TEM grid with thin-section imaged
worm_id sperm_id - worm from which thin-section stemmed
magnification - total magnification at which an image was taken (this is used to translate image area into biological area using ImageJ)
sperm_surfacearea_total - total area of the sperm cell measured using ImageJ
MO_surfacearea_1 - area of first-measured membranous organelle (MO) of the sperm cell measured using ImageJ (the order in which MOs within a sperm cell are measured is arbitrary and columns recording various MOs are numbered only for data analysis purposes)
MO_surfacearea_2 - area of second membranous organelle (MO) of the sperm cell measured using ImageJ 
MO_surfacearea_3 - area of third membranous organelle (MO) of the sperm cell measured using ImageJ 
MO_surfacearea_4 - area of fourth membranous organelle (MO) of the sperm cell measured using ImageJ 
MO_surfacearea_5 - area of fifth membranous organelle (MO) of the sperm cell measured using ImageJ 
MO_surfacearea_6 - area of sixth membranous organelle (MO) of the sperm cell measured using ImageJ
MO_surfacearea_7 - area of seventh membranous organelle (MO) of the sperm cell measured using ImageJ 
MO_surfacearea_8 - area of eight membranous organelle (MO) of the sperm cell measured using ImageJ 
MO_surfacearea_9 - area of nineth membranous organelle (MO) of the sperm cell measured using ImageJ 
MO_surfacearea_10 - area of tenth membranous organelle (MO) of the sperm cell measured using ImageJ 
MO_surfacearea_11 - area of 11th membranous organelle (MO) of the sperm cell measured using ImageJ 
MO_surfacearea_12 - area of 12th membranous organelle (MO) of the sperm cell measured using ImageJ 
MO_surfacearea_13 - area of 13th membranous organelle (MO) of the sperm cell measured using ImageJ 
MO_surfacearea_14 - area of 14th membranous organelle (MO) of the sperm cell measured using ImageJ 
MO_surfacearea_15 - area of 15th membranous organelle (MO) of the sperm cell measured using ImageJ 
MO_surfacearea_16 - area of 16th membranous organelle (MO) of the sperm cell measured using ImageJ 
MO_surfacearea_17 - area of 17th membranous organelle (MO) of the sperm cell measured using ImageJ 
MO_surfacearea_18 - area of 18th membranous organelle (MO) of the sperm cell measured using ImageJ 
MO_surfacearea_19 - area of 19th membranous organelle (MO) of the sperm cell measured using ImageJ 
MO_surfacearea_20 - area of 20th membranous organelle (MO) of the sperm cell measured using ImageJ 
MO_surfacearea_21 - area of 21st membranous organelle (MO) of the sperm cell measured using ImageJ 
MO_surfacearea_22 - area of 22nd membranous organelle (MO) of the sperm cell measured using ImageJ 
MO_surfacearea_23 - area of 23rd membranous organelle (MO) of the sperm cell measured using ImageJ 
MO_surfacearea_24 - area of 24th membranous organelle (MO) of the sperm cell measured using ImageJ 
MO_surfacearea_25 - area of 25th membranous organelle (MO) of the sperm cell measured using ImageJ 
MO_surfacearea_26 - area of 26th membranous organelle (MO) of the sperm cell measured using ImageJ 
MO_surfacearea_27 - area of 27th membranous organelle (MO) of the sperm cell measured using ImageJ 
MO_surfacearea_28 - area of 28th membranous organelle (MO) of the sperm cell measured using ImageJ 
MO_surfacearea_29 - area of 29th membranous organelle (MO) of the sperm cell measured using ImageJ 
MO_surfacearea_30 - area of 30th membranous organelle (MO) of the sperm cell measured using ImageJ 
mitochondrion_surfacearea_1 - area of first-measured mitochondrion of the sperm cell measured using ImageJ (the order in which mitochondria within a sperm cell are measured is arbitrary and columns recording various mitochondria are numbered only for data analysis purposes)
mitochondrion_surfacearea_2 - area of second mitochondrion of the sperm cell measured using ImageJ 
mitochondrion_surfacearea_3 - area of third mitochondrion of the sperm cell measured using ImageJ
mitochondrion_surfacearea_4 - area of fourth mitochondrion of the sperm cell measured using ImageJ
mitochondrion_surfacearea_5 - area of fifth mitochondrion of the sperm cell measured using ImageJ
mitochondrion_surfacearea_6 - area of sixth mitochondrion of the sperm cell measured using ImageJ
mitochondrion_surfacearea_7 - area of seventh mitochondrion of the sperm cell measured using ImageJ
mitochondrion_surfacearea_8 - area of eighth mitochondrion of the sperm cell measured using ImageJ
mitochondrion_surfacearea_9 - area of nineth mitochondrion of the sperm cell measured using ImageJ
mitochondrion_surfacearea_10 - area of tenth mitochondrion of the sperm cell measured using ImageJ
mitochondrion_surfacearea_11 - area of 11th mitochondrion of the sperm cell measured using ImageJ
mitochondrion_surfacearea_12 - area of 12th mitochondrion of the sperm cell measured using ImageJ
mitochondrion_surfacearea_13 - area of 13th mitochondrion of the sperm cell measured using ImageJ
mitochondrion_surfacearea_14 - area of 14th mitochondrion of the sperm cell measured using ImageJ
mitochondrion_surfacearea_15 - area of 15th mitochondrion of the sperm cell measured using ImageJ
mitochondrion_surfacearea_16 - area of 16th mitochondrion of the sperm cell measured using ImageJ
mitochondrion_surfacearea_17 - area of 17th mitochondrion of the sperm cell measured using ImageJ
mitochondrion_surfacearea_18 - area of 18th mitochondrion of the sperm cell measured using ImageJ
mitochondrion_surfacearea_19 - area of 19th mitochondrion of the sperm cell measured using ImageJ
mitochondrion_surfacearea_20 - area of 20th mitochondrion of the sperm cell measured using ImageJ
mitochondrion_surfacearea_21 - area of 21st mitochondrion of the sperm cell measured using ImageJ
mitochondrion_surfacearea_22 - area of 22nd mitochondrion of the sperm cell measured using ImageJ
mitochondrion_surfacearea_23 - area of 23rd mitochondrion of the sperm cell measured using ImageJ
mitochondrion_surfacearea_24 - area of 24th mitochondrion of the sperm cell measured using ImageJ
mitochondrion_surfacearea_25 - area of 25th mitochondrion of the sperm cell measured using ImageJ
mitochondrion_surfacearea_26 - area of 26th mitochondrion of the sperm cell measured using ImageJ
mitochondrion_surfacearea_27 - area of 27th mitochondrion of the sperm cell measured using ImageJ
mitochondrion_surfacearea_28 - area of 28th mitochondrion of the sperm cell measured using ImageJ
mitochondrion_surfacearea_29 - area of 29th mitochondrion of the sperm cell measured using ImageJ
mitochondrion_surfacearea_30 - area of 30th mitochondrion of the sperm cell measured using ImageJ
mitochondrion_surfacearea_31 - area of 31st mitochondrion of the sperm cell measured using ImageJ
nucleus_surfacearea - if a nucleus was detected in a given cell's cross section, it's area was measured and recorded (darkest area of the cell) using ImageJ
pseudopod_surfacearea - area measurement of granulous, non-cell body area with MSP using ImageJ
date_embedded_yearmonthday - date the worm in this sample was fixed and resin-embedded (for internal tracking of physical samples)
censornomito - column for excluding sperm cells with zero recorded mitochondria (no mitochondria suggests the sperm cell was sectioned too close to the cell wall for analysis)
measuredby - person who recorded the area of a sperm cell using Image J
imagequality.1.best_3.worst. - qualitative metric for recording if a specific samples was of poor quality, possibly affecting confidence in measurements

**Note: any values reported in this table correspond to raw data measured by Christine Rehaluk using ImageJ. All wrangling and processing of data occurs in the accompanying TEMms.Rmd file

2 - TEMms.Rmd is the script used to wrangle data and create data figures for the manuscript
