## Supplementary material for "Divergent organelle allocation in the evolution of sperm gigantism revealed from subcellular quantification of nematode sperm with electron microscopy": data and analysis scripts: TEMms.Rmd

````
TEM sperm cell and organelle areas in C. nouraguensis and C. macrosperma:

The purpose of this study is to explore differences between sperm cell composition in species with giant vs. standard sized sperm. To accomplish this, mated females with sperm in their uteri of the species C. nouraguensis (standard-sized sperm) and C. macrosperma (giant sperm) were chemically fixed, embedded, thin-sectioned, stained, and TEM imaged. This allows us to view cross-sections of sperm cells inside the uterus and, using ImageJ software, measure the area of the total sperm cell, pseudopod, and organelles such as mitochondria, membranous organelles (MOs), and in some cases nuclei. Additionally, we can compare relative area of a sperm cell allocated to an organelle type as a proxy for certain investment into that organelle and its function in the sperm cell.

Set-up: working directory, packages, load data
```{r Load Packages & Data, message=FALSE, warning=FALSE}
# Load all the data and all the packages for whole script
library(tidyverse)
library(ggplot2)
library(ggrepel)
library(RColorBrewer)
library(svglite)
library(gridExtra)
library(cowplot)

getwd()

alltemdata<-read.csv("data/SpermCellAreaDataRecordedValuesOnly.csv") #TEM image analysis data with cell and organelle area data

```

Checking in to make sure all data is there and read correctly, including any necessary data wrangling
```{r Check Data, message=FALSE, warning=FALSE}
#Check data
head(alltemdata)
str(alltemdata)

```

Count how many instances there are of each organelle (MO and mitochondria) per sperm cell (=organelle count)
```{r Organelle Counts, message=FALSE, warning=FALSE}
# Estimate number of MOs per cell
names(alltemdata)[9:38]  ## the columns with the MO info
alltemdata$MO_count <- sapply(1:nrow(alltemdata), function(x) sum(!is.na(alltemdata[x, 9:38])))  ## create the MO count column
  
# Estimate number of mitochondria per cell
names(alltemdata)[39:69]  ## same for mito count
alltemdata$mitochondria_count <- sapply(1:nrow(alltemdata), function(x) sum(!is.na(alltemdata[x, 39:69])))

```

Censor data from replicate sperm cells without mitochondria in them (thin-section not far enough into the cell body to be informative)
```{r Censor replicates without mitochondria, message=FALSE, warning=FALSE}
temdata <- subset(alltemdata, alltemdata$mitochondria_count != 0) # censor all replicates without mitochondria
temdata$species = as.factor(temdata$species)

head(temdata)
str(temdata)

```

Measure length of column entries per species to have replicate number after censoring replicates without mitochondria
```{r Retrieve replication per species, message=FALSE, warning=FALSE}
## variable with number of replicates
n_Cmacro <- length(temdata$species[temdata$species == "C. macrosperma"])
n_Cnoura <- length(temdata$species[temdata$species ==  "C. nouraguensis"])

```

Make columns for various variables we are looking to plot. Start by summing up total areas for organelles with multiple instances per sperm cell.
```{r Totals of organelle areas, message=FALSE, warning=FALSE}
# Estimate total area of MOs
temdata$MO_surfacearea_total <- rowSums(temdata[,c("MO_surfacearea_1", "MO_surfacearea_2", "MO_surfacearea_3", "MO_surfacearea_4", "MO_surfacearea_5", "MO_surfacearea_6", "MO_surfacearea_7", "MO_surfacearea_8", "MO_surfacearea_9", "MO_surfacearea_10", "MO_surfacearea_11", "MO_surfacearea_12", "MO_surfacearea_13", "MO_surfacearea_14", "MO_surfacearea_15", "MO_surfacearea_16", "MO_surfacearea_17", "MO_surfacearea_18", "MO_surfacearea_19", "MO_surfacearea_20", "MO_surfacearea_21", "MO_surfacearea_22", "MO_surfacearea_23", "MO_surfacearea_24", "MO_surfacearea_25", "MO_surfacearea_26", "MO_surfacearea_27", "MO_surfacearea_28", "MO_surfacearea_29", "MO_surfacearea_30")], na.rm=TRUE)

# Estimate total area of mitochondria
temdata$mitochondria_surfacearea_total <- rowSums(temdata[,c("mitochondrion_surfacearea_1", "mitochondrion_surfacearea_2", "mitochondrion_surfacearea_3", "mitochondrion_surfacearea_4", "mitochondrion_surfacearea_5", "mitochondrion_surfacearea_6", "mitochondrion_surfacearea_7", "mitochondrion_surfacearea_8", "mitochondrion_surfacearea_9", "mitochondrion_surfacearea_10", "mitochondrion_surfacearea_11", "mitochondrion_surfacearea_12", "mitochondrion_surfacearea_13", "mitochondrion_surfacearea_14", "mitochondrion_surfacearea_15", "mitochondrion_surfacearea_16", "mitochondrion_surfacearea_17", "mitochondrion_surfacearea_18", "mitochondrion_surfacearea_19", "mitochondrion_surfacearea_20", "mitochondrion_surfacearea_21", "mitochondrion_surfacearea_22", "mitochondrion_surfacearea_23", "mitochondrion_surfacearea_24", "mitochondrion_surfacearea_25", "mitochondrion_surfacearea_26", "mitochondrion_surfacearea_27", "mitochondrion_surfacearea_28", "mitochondrion_surfacearea_29", "mitochondrion_surfacearea_30", "mitochondrion_surfacearea_31")], na.rm=TRUE)
```

Estimate other cell areas from measured components
```{r Cellbody and "other" cell areas, message=FALSE, warning=FALSE}
# Estimate cell plasma and other unidentifiable area, by taking total cell size and subtracting all identified organelles and the pseudopod
temdata$other_surfacearea <- temdata$sperm_surfacearea_total - temdata$MO_surfacearea_total - temdata$mitochondria_surfacearea_total - temdata$nucleus_surfacearea - temdata$pseudopod_surfacearea

# Estimate area of cell body only, without pseudopod
temdata$cellbodywithoutpseudopod <- temdata$sperm_surfacearea_total - temdata$pseudopod_surfacearea

```

Standardize organelle area by total area, for whole cell, as well as cell body only. Also estimate fraction of overall cell that is pseudopod.
```{r Organelles & pseudopod per total cell and cell body, message=FALSE, warning=FALSE}
# Standardize total area of mitochondria for whole cell (cell body and pseudopod)
temdata$mitopertotalcell <- temdata$mitochondria_surfacearea_total / temdata$sperm_surfacearea_total
  
# Standardize total area of MOs for whole cell (cell body and pseudopod)
temdata$MOpertotalcell <- temdata$MO_surfacearea_total / temdata$sperm_surfacearea_total

# Standardize total area of mitochondria for cell body only (without pseudopod)
temdata$mitopercellbody <- temdata$mitochondria_surfacearea_total / temdata$cellbodywithoutpseudopod
  
# Standardize total area of MOs for cell body only (without pseudopod)
temdata$MOpercellbody <- temdata$MO_surfacearea_total / temdata$cellbodywithoutpseudopod

# Estimate fraction of pseudopod per overall cell size
temdata$pseudopodpertotalcell <- temdata$pseudopod_surfacearea / temdata$sperm_surfacearea_total

```

Estimate average size (area) of mitochondria and MO
```{r Organelle average sizes (area), message=FALSE, warning=FALSE}
# Estimate average size of an MO-body in a cell
temdata$MO_average <- temdata$MO_surfacearea_total/temdata$MO_count

# Estimate average size of mitochondrion per cell
temdata$mitochondria_average <- temdata$mitochondria_surfacearea_total/temdata$mitochondria_count

```

Make columns estimating idealized sphere
```{r Idealized sphere circumference and MOs, message=FALSE, warning=FALSE}

## Generate idealized cell circumference from measure area -> idealized radius -> idealized circumference

## area: a = pi*r^2   so r = sqrt(a/pi)
## circumference: c = 2*pi*r  so c = 2*pi*sqrt(a/pi)
GetCircumfrenceFromArea<-function(a) 2*pi*sqrt(a/pi)

## Make column with circumference of idealized circle from sperm cell BODY area [um]
temdata$IdealizedCircumference = sapply(1:nrow(temdata), function(x) GetCircumfrenceFromArea(temdata$cellbodywithoutpseudopod [x]))

## Standardize MO-bodies per idealized circumference
temdata$MOpercircumference <- temdata$MO_count/temdata$IdealizedCircumference

## Standardize MO-body area per idealized circumference
temdata$MOareapercircumference <- temdata$MO_surfacearea_total/temdata$IdealizedCircumference

```

Generate density variable for mitochondria
```{r Mitochondria count per cell area, message=FALSE, warning=FALSE}
## Mitochondrial density as mitochondria per cell body area
temdata$mitodensity <- temdata$mitochondria_count / temdata$cellbodywithoutpseudopod

```

Estimate MO-area for total ejaculates
```{r MO per ejaculate, message=FALSE, warning=FALSE}

macroejaculate <- 230 # approximated from Vielle et al. 2016
nouraejaculate <- 475 # approximated from Vielle et al. 2016

## Estimate average MO area per whole ejaculate
#MOperMacroEjac <- mean(temdata[temdata$species == "C. macrosperma", "MO_surfacearea_total"])*macroejaculate
#MOperMacroEjac 

#MOperNouraEjac <- mean(temdata[temdata$species == "C. nouraguensis", "MO_surfacearea_total"])*nouraejaculate
#MOperNouraEjac

## Estimate average MO volume from average MO area
# Average MO area from mean of Average MO size 
averageMOareaMacro <- mean(temdata[temdata$species == "C. macrosperma", "MO_average"])
averageMOareaMacro

averageMOareaNoura <- mean(temdata[temdata$species == "C. nouraguensis", "MO_average"])
averageMOareaNoura

# Retrieve radius from area of idealzed circle of the same area as our measured MOs
radiusAverageMOmacro <- sqrt(averageMOareaMacro/pi)
radiusAverageMOmacro

radiusAverageMOnoura <- sqrt(averageMOareaNoura/pi)
radiusAverageMOnoura

# Estimate volume of this idealized sphere (um^3) that shares its cross-sectional area with the average sized MO of either species using V=4/3*pi*r^3
volumeAverageMOmacro <- (4/3)*pi*radiusAverageMOmacro^3
volumeAverageMOmacro

volumeAverageMOnoura <- (4/3)*pi*radiusAverageMOnoura^3
volumeAverageMOnoura

# Estimate collective volume of a species ejaculate (um^3) by multiplying the volume of an average MO with the average number of MOs per cell and the number of cells per ejaculate (inferred from Vielle et al., 2016)
MacroMOvolumeperejaculate <- volumeAverageMOmacro*mean(temdata[temdata$species == "C. macrosperma", "MO_count"])*macroejaculate
MacroMOvolumeperejaculate

NouraMOvolumeperejaculate <- volumeAverageMOnoura*mean(temdata[temdata$species == "C. nouraguensis", "MO_count"])*nouraejaculate
NouraMOvolumeperejaculate

```

Pull up table with all the headers that were newly created
```{r View data table, message=FALSE, warning=FALSE}
 View(temdata)
```

Plots for final manuscript
```{r Final plots & t-test, message=FALSE, warning=FALSE}

# find identifier to determine species order in all graphs
unique(temdata$species_name)
# determine order
temdata$species = factor(temdata$species, levels=c(
  "C. nouraguensis", "C. macrosperma"
))

# Assign colours for each species
cols2 = c("#7E7E7E", "#212121")

# Create theme for all box plots
TEMtheme <-   
  theme(panel.background = element_rect(fill = "transparent"),
        panel.grid.minor = element_blank(),
        panel.grid.major = element_blank())+
  theme(plot.background = element_rect(fill = "transparent"),
        legend.background = element_rect(fill = "transparent"),
        legend.text = element_text(colour = "black", size = 12),
        legend.title = element_text(colour = "black", size = 15),
        legend.key = element_rect(fill = "transparent")) + 
  theme(legend.position = "none") +
  theme(axis.text.x = element_text(face='italic', colour = "black", size = 12), 
        axis.text.y = element_text(colour = "black", size = 12),
        axis.line = element_line(colour="black", linetype = "solid"),
        axis.ticks = element_line(colour="black", linetype = "solid"),
        axis.title.x = element_text(colour = "black", size = 15),
        axis.title.y = element_text(margin=margin(l=10), colour = "black", size = 15))
####

# Figure 1: sperm cell schematic and TEM images of spermatozoa
## see manuscript

# Figure 2: total sperm cell size, cell body, total pseudopod, standardized pseudopod

# Figure 2A: Total sperm cell area in µm^2
q2A.box = ggplot(temdata, aes(species, sperm_surfacearea_total, color=species)) + 
  geom_boxplot(alpha = 0, lwd = 1) + 
  geom_jitter(alpha = 0.55, width = 0.2, size = 2) +
  labs(x = " ", y = bquote("Total sperm cell area "~(µm^2))) +
  scale_color_manual(values=cols2, name="species") +
  TEMtheme
q2A.box 
# save as pdf
ggsave(q2A.box, width=4, height=4, dpi = 600, units=c("in"), filename="output/Figure2A_TotalSpermCellArea.pdf", bg="transparent")

# test differences in total sperm size (with pseudopod) (report in table: t-statistic, df, p value, confidence?, mean of x and mean of y, SE of difference)
t.test(temdata[temdata$species == "C. nouraguensis", "sperm_surfacearea_total"], temdata[temdata$species == "C. macrosperma", "sperm_surfacearea_total"], paired = F)

SEofCmacroTotalSpermCell <- (sd(temdata[temdata$species == "C. macrosperma", "sperm_surfacearea_total"]))/sqrt(n_Cmacro)
SEofCmacroTotalSpermCell

SEofCnouraTotalSpermCell <- (sd(temdata[temdata$species == "C. nouraguensis", "sperm_surfacearea_total"]))/sqrt(n_Cnoura)
SEofCnouraTotalSpermCell
####

# Figure 2B: Sperm cell body area only in µm^2 (no psudopod)
q2B.box = ggplot(temdata, aes(species, cellbodywithoutpseudopod, color=species)) + 
  geom_boxplot(alpha = 0, lwd = 1) + 
  geom_jitter(alpha = 0.55, width = 0.2, size = 2) +
  labs(x = " ", y = "Sperm cell body area "~(µm^2)) +
  scale_color_manual(values=cols2, name="species") +
  TEMtheme
q2B.box 
# save as pdf
ggsave(q2B.box, width=4, height=4, dpi = 600, units=c("in"), filename="output/Figure2B_SpermCellBodyOnlyArea.pdf", bg="transparent")

# test differences in sperm size (without pseudopod) (report in table: t-statistic, df, p value, confidence?, mean of x and mean of y, SE of difference)
t.test(temdata[temdata$species == "C. nouraguensis", "cellbodywithoutpseudopod"], temdata[temdata$species == "C. macrosperma", "cellbodywithoutpseudopod"], paired = F)

SEofCmacroSpermCellBody <- (sd(temdata[temdata$species == "C. macrosperma", "cellbodywithoutpseudopod"]))/sqrt(n_Cmacro)
SEofCmacroSpermCellBody

SEofCnouraSpermCellBody <- (sd(temdata[temdata$species == "C. nouraguensis", "cellbodywithoutpseudopod"]))/sqrt(n_Cnoura)
SEofCnouraSpermCellBody
####

# Figure 2C: Total pseudopod area in µm^2
q2C.box = ggplot(temdata, aes(species, pseudopod_surfacearea, color=species)) + 
  geom_boxplot(alpha = 0, lwd = 1) + 
  geom_jitter(alpha = 0.55, width = 0.2, size = 2) +
  labs(x = " ", y = "Total pseudopod area "~(µm^2)) +
  scale_color_manual(values=cols2, name="species") +
  TEMtheme
q2C.box 
# save as pdf
ggsave(q2C.box, width=4, height=4, dpi = 600, units=c("in"), filename="output/Figure2C_TotalPseudopodArea.pdf", bg="transparent")

# test differences in total pseudopod area (report in table: t-statistic, df, p value, confidence?, mean of x and mean of y, SE of difference)
t.test(temdata[temdata$species == "C. nouraguensis", "pseudopod_surfacearea"], temdata[temdata$species == "C. macrosperma", "pseudopod_surfacearea"], paired = F)

SEofCmacroFalsebean<- (sd(temdata[temdata$species == "C. macrosperma", "pseudopod_surfacearea"]))/sqrt(n_Cmacro)
SEofCmacroFalsebean

SEofCnouraFalsebean <- (sd(temdata[temdata$species == "C. nouraguensis", "pseudopod_surfacearea"]))/sqrt(n_Cnoura)
SEofCnouraFalsebean
####

# Figure 2D: Standardized area of pseudopod per total cell in µm^2
q2D.box = ggplot(temdata, aes(species, pseudopodpertotalcell, color=species)) + 
  geom_boxplot(alpha = 0, lwd = 1) + 
  geom_jitter(alpha = 0.55, width = 0.2, size = 2) +
  labs(x = " ", y = "Relative pseudopod area") +
  scale_color_manual(values=cols2, name="species") +
  TEMtheme
q2D.box 
# save as pdf
ggsave(q2D.box, width=4, height=4, dpi = 600, units=c("in"), filename="output/Figure2D_StandardizedPseudopodAreaPerSpermCell.pdf", bg="transparent")

# test differences in pseudopod area per total sperm cell (report in table: t-statistic, df, p value, confidence?, mean of x and mean of y, SE of difference)
t.test(temdata[temdata$species == "C. nouraguensis", "pseudopodpertotalcell"], temdata[temdata$species == "C. macrosperma", "pseudopodpertotalcell"], paired = F)

SEofCmacroStandardizedFalsebean<- (sd(temdata[temdata$species == "C. macrosperma", "pseudopodpertotalcell"]))/sqrt(n_Cmacro)
SEofCmacroStandardizedFalsebean

SEofCnouraStandardizedFalsebean <- (sd(temdata[temdata$species == "C. nouraguensis", "pseudopodpertotalcell"]))/sqrt(n_Cnoura)
SEofCnouraStandardizedFalsebean
####

# Assemble 4 panels and add p-values
q1 <- q2A.box + theme(legend.position = "none")  + theme(axis.title.x = element_blank())
q1 <- ggdraw(q1) + draw_plot_label(label = "A",x = 0, y = 1, hjust = -2, vjust = 2, colour = "black", size = 16, fontface = "bold")
q2 <- q2B.box + theme(legend.position = "none")  + theme(axis.title.x = element_blank())
q2 <- ggdraw(q2) + draw_plot_label(label = "B",x = 0, y = 1, hjust = -2, vjust = 2, colour = "black", size = 16, fontface = "bold")
q3 <- q2C.box + theme(legend.position = "none")  + theme(axis.title.x = element_blank())
q3 <- ggdraw(q3) + draw_plot_label(label = "C",x = 0, y = 1, hjust = -2, vjust = 2, colour = "black", size = 16, fontface = "bold")
q4 <- q2D.box + theme(legend.position = "none") + theme(axis.title.x = element_blank())
q4 <- ggdraw(q4) + draw_plot_label(label = "D",x = 0, y = 1, hjust = -2, vjust = 2, colour = "black", size = 16, fontface = "bold")

qplot <- grid.arrange(q1, q2, q3, q4, ncol = 2, nrow = 2)
qplot
ggsave(qplot, width=25, height=25, dpi = 600, units=c("cm"), filename="output/Figure2_CellSize&Pseudopods.pdf", bg="transparent")
####

# Figure 3: Mitochondria

# Figure 3A: Total number of mitochondria per cell
q3A.box = ggplot(temdata, aes(species, mitochondria_count, color=species)) + 
  geom_boxplot(alpha = 0, lwd = 1) + 
  geom_jitter(alpha = 0.55, width = 0.2, size = 2) +
  labs(x = " ", y = "Total number of mitochondria") +
  scale_color_manual(values=cols2, name="species") +
  TEMtheme
q3A.box 
# save as pdf
ggsave(q3A.box, width=4, height=4, dpi = 600, units=c("in"), filename="output/Figure3A_TotalNumberofMitochondria.pdf", bg="transparent")

# test differences in total mitochondria count (report in table: t-statistic, df, p value, confidence?, mean of x and mean of y, SE of difference)
t.test(temdata[temdata$species == "C. nouraguensis", "mitochondria_count"], temdata[temdata$species == "C. macrosperma", "mitochondria_count"], paired = F)

SEofCmacroMitocount<- (sd(temdata[temdata$species == "C. macrosperma", "mitochondria_count"]))/sqrt(n_Cmacro)
SEofCmacroMitocount

SEofCnouraMitocount <- (sd(temdata[temdata$species == "C. nouraguensis", "mitochondria_count"]))/sqrt(n_Cnoura)
SEofCnouraMitocount
####

# Figure 3B: Average area of mitochondrion in a cell in µm^2
q3B.box = ggplot(temdata, aes(species, mitochondria_average, color=species)) + 
  geom_boxplot(alpha = 0, lwd = 1) + 
  geom_jitter(alpha = 0.55, width = 0.2, size = 2) +
  labs(x = " ", y = "Average area of \na mitochondrion "~(µm^2)) +
  scale_color_manual(values=cols2, name="species") +
  TEMtheme
q3B.box 
# save as pdf
ggsave(q3B.box, width=4.2, height=4, dpi = 600, units=c("in"), filename="output/Figure3B_AverageMitochondrion.pdf", bg="transparent")

# test differences in average mitochondrial area (report in table: t-statistic, df, p value, confidence?, mean of x and mean of y, SE of difference)
t.test(temdata[temdata$species == "C. nouraguensis", "mitochondria_average"], temdata[temdata$species == "C. macrosperma", "mitochondria_average"], paired = F)

SEofCmacroMitoaverage<- (sd(temdata[temdata$species == "C. macrosperma", "mitochondria_average"]))/sqrt(n_Cmacro)
SEofCmacroMitoaverage

SEofCnouraMitoaverage <- (sd(temdata[temdata$species == "C. nouraguensis", "mitochondria_average"]))/sqrt(n_Cnoura)
SEofCnouraMitoaverage
####

# Figure 3C: Total mitochondrial area in µm^2
q3C.box = ggplot(temdata, aes(species, mitochondria_surfacearea_total, color=species)) + 
  geom_boxplot(alpha = 0, lwd = 1) + 
  geom_jitter(alpha = 0.55, width = 0.2, size = 2) +
  labs(x = " ", y = "Total mitochondrial area "~(µm^2)) +
  scale_color_manual(values=cols2, name="species") +
  TEMtheme
q3C.box 
# save as pdf
ggsave(q3C.box, width=4, height=4, dpi = 600, units=c("in"), filename="output/Figure3C_TotalMitochondrialArea.pdf", bg="transparent")

# test differences in total mitochondrial area (report in table: t-statistic, df, p value, confidence?, mean of x and mean of y, SE of difference)
t.test(temdata[temdata$species == "C. nouraguensis", "mitochondria_surfacearea_total"], temdata[temdata$species == "C. macrosperma", "mitochondria_surfacearea_total"], paired = F)

SEofCmacroTotalMito<- (sd(temdata[temdata$species == "C. macrosperma", "mitochondria_surfacearea_total"]))/sqrt(n_Cmacro)
SEofCmacroTotalMito

SEofCnouraTotalMito <- (sd(temdata[temdata$species == "C. nouraguensis", "mitochondria_surfacearea_total"]))/sqrt(n_Cnoura)
SEofCnouraTotalMito
####

# Figure 3D: Standardized area of mitochondria per cell body in µm^2
q3D.box = ggplot(temdata, aes(species, mitopercellbody, color=species)) + 
  geom_boxplot(alpha = 0, lwd = 1) + 
  geom_jitter(alpha = 0.55, width = 0.2, size = 2) +
  labs(x = " ", y = "Relative mitochondrial area \nper sperm cell body") +
  scale_color_manual(values=cols2, name="species") +
  TEMtheme
q3D.box 
# save as pdf
ggsave(q3D.box, width=4.2, height=4, dpi = 600, units=c("in"), filename="output/Figure3D_StandardizedMitochondriaAreaPerCellBody.pdf", bg="transparent")

# test differences in standardized mitochondria area per cell body only (no pseudopod) (report in table: t-statistic, df, p value, confidence?, mean of x and mean of y, SE of difference)
t.test(temdata[temdata$species == "C. nouraguensis", "mitopercellbody"], temdata[temdata$species == "C. macrosperma", "mitopercellbody"], paired = F)

SEofCmacroMitopercellbody<- (sd(temdata[temdata$species == "C. macrosperma", "mitopercellbody"]))/sqrt(n_Cmacro)
SEofCmacroMitopercellbody

SEofCnouraMitopercellbody <- (sd(temdata[temdata$species == "C. nouraguensis", "mitopercellbody"]))/sqrt(n_Cnoura)
SEofCnouraMitopercellbody
####

# Assemble 4 panels and add p-values
q1 <- q3A.box + theme(legend.position = "none")  + theme(axis.title.x = element_blank())
q1 <- ggdraw(q1) + draw_plot_label(label = "A",x = 0, y = 1, hjust = -2, vjust = 2, colour = "black", size = 16, fontface = "bold")
q2 <- q3B.box + theme(legend.position = "none")  + theme(axis.title.x = element_blank())
q2 <- ggdraw(q2) + draw_plot_label(label = "B",x = 0, y = 1, hjust = -2, vjust = 2, colour = "black", size = 16, fontface = "bold")
q3 <- q3C.box + theme(legend.position = "none")  + theme(axis.title.x = element_blank())
q3 <- ggdraw(q3) + draw_plot_label(label = "C",x = 0, y = 1, hjust = -2, vjust = 2, colour = "black", size = 16, fontface = "bold")
q4 <- q3D.box + theme(legend.position = "none") + theme(axis.title.x = element_blank())
q4 <- ggdraw(q4) + draw_plot_label(label = "D",x = 0, y = 1, hjust = -2, vjust = 2, colour = "black", size = 16, fontface = "bold")

qplot <- grid.arrange(q1, q2, q3, q4, ncol = 2, nrow = 2)
qplot
ggsave(qplot, width=25, height=25, dpi = 600, units=c("cm"), filename="output/Figure3_Mitochondria.pdf", bg="transparent")
####

# Figure 4: MOs

# Figure 4A: Total number of MOs per cell
q4A.box = ggplot(temdata, aes(species, MO_count, color=species)) + 
  geom_boxplot(alpha = 0, lwd = 1) + 
  geom_jitter(alpha = 0.55, width = 0.2, size = 2) +
  labs(x = " ", y = "Total number of MOs") +
  scale_color_manual(values=cols2, name="species") +
  TEMtheme
q4A.box 
# save as pdf
ggsave(q4A.box, width=4, height=4, dpi = 600, units=c("in"), filename="output/Figure4A_TotalNumberofMOs.pdf", bg="transparent")

# test differences in total MO count (report in table: t-statistic, df, p value, confidence?, mean of x and mean of y, SE of difference)
t.test(temdata[temdata$species == "C. nouraguensis", "MO_count"], temdata[temdata$species == "C. macrosperma", "MO_count"], paired = F)

SEofCmacroMOcount<- (sd(temdata[temdata$species == "C. macrosperma", "MO_count"]))/sqrt(n_Cmacro)
SEofCmacroMOcount

SEofCnouraMOcount <- (sd(temdata[temdata$species == "C. nouraguensis", "MO_count"]))/sqrt(n_Cnoura)
SEofCnouraMOcount
####

# Figure 4B: Average area of MO in a cell in µm^2
q4B.box = ggplot(temdata, aes(species, MO_average, color=species)) + 
  geom_boxplot(alpha = 0, lwd = 1) + 
  geom_jitter(alpha = 0.55, width = 0.2, size = 2) +
  labs(x = " ", y = "Average area of an MO "~(µm^2)) +
  scale_color_manual(values=cols2, name="species") +
  TEMtheme
q4B.box 
# save as pdf
ggsave(q4B.box, width=4, height=4, dpi = 600, units=c("in"), filename="output/Figure4B_AverageMOAreaPerSpermCell.pdf", bg="transparent")

# test differences in average MO area (report in table: t-statistic, df, p value, confidence?, mean of x and mean of y, SE of difference)
t.test(temdata[temdata$species == "C. nouraguensis", "MO_average"], temdata[temdata$species == "C. macrosperma", "MO_average"], paired = F)

SEofCmacroMOaverage<- (sd(temdata[temdata$species == "C. macrosperma", "MO_average"]))/sqrt(n_Cmacro)
SEofCmacroMOaverage

SEofCnouraMOaverage <- (sd(temdata[temdata$species == "C. nouraguensis", "MO_average"]))/sqrt(n_Cnoura)
SEofCnouraMOaverage
####

# Figure 4C: Total MO area in µm^2
q4C.box = ggplot(temdata, aes(species, MO_surfacearea_total, color=species)) + 
  geom_boxplot(alpha = 0, lwd = 1) + 
  geom_jitter(alpha = 0.55, width = 0.2, size = 2) +
  labs(x = " ", y = "Total MO area "~(µm^2)) +
  scale_color_manual(values=cols2, name="species") +
  TEMtheme
q4C.box 
# save as pdf
ggsave(q4C.box, width=4, height=4, dpi = 600, units=c("in"), filename="output/Figure4C_TotalMOArea.pdf", bg="transparent")

# test differences in total MO area (report in table: t-statistic, df, p value, confidence?, mean of x and mean of y, SE of difference)
t.test(temdata[temdata$species == "C. nouraguensis", "MO_surfacearea_total"], temdata[temdata$species == "C. macrosperma", "MO_surfacearea_total"], paired = F)

SEofCmacroTotalMO<- (sd(temdata[temdata$species == "C. macrosperma", "MO_surfacearea_total"]))/sqrt(n_Cmacro)
SEofCmacroTotalMO

SEofCnouraTotalMO <- (sd(temdata[temdata$species == "C. nouraguensis", "MO_surfacearea_total"]))/sqrt(n_Cnoura)
SEofCnouraTotalMO
####

# Figure 4D: MO area per idealized cell body circumference or Peripheral MO area
q4D.box = ggplot(temdata, aes(species, MOareapercircumference, color=species)) + 
  geom_boxplot(alpha = 0, lwd = 1) + 
  geom_jitter(alpha = 0.55, width = 0.2, size = 2) +
  labs(x = " ", y = "Peripheral MO area "~(µm^2/µm)) +
  scale_color_manual(values=cols2, name="species") +
  TEMtheme
q4D.box 
# save as pdf
ggsave(q4D.box, width=4.2, height=4, dpi = 600, units=c("in"), filename="output/Figure4D_AreaOfMOsPerCellBodyCircumference.pdf", bg="transparent")

# test differences in area of MO-bodies area per idealized cell body circumference (report in table: t-statistic, df, p value, confidence?, mean of x and mean of y, SE of difference)
t.test(temdata[temdata$species == "C. nouraguensis", "MOareapercircumference"], temdata[temdata$species == "C. macrosperma", "MOareapercircumference"], paired = F)

SEofCmacroMOareapercircumference<- (sd(temdata[temdata$species == "C. macrosperma", "MOareapercircumference"]))/sqrt(n_Cmacro)
SEofCmacroMOareapercircumference

SEofCnouraMOareapercircumference <- (sd(temdata[temdata$species == "C. nouraguensis", "MOareapercircumference"]))/sqrt(n_Cnoura)
SEofCnouraMOareapercircumference
####

# Assemble 4 panels and add p-values
q1 <- q4A.box + theme(legend.position = "none")  + theme(axis.title.x = element_blank())
q1 <- ggdraw(q1) + draw_plot_label(label = "A",x = 0, y = 1, hjust = -2, vjust = 2, colour = "black", size = 16, fontface = "bold")
q2 <- q4B.box + theme(legend.position = "none")  + theme(axis.title.x = element_blank())
q2 <- ggdraw(q2) + draw_plot_label(label = "B",x = 0, y = 1, hjust = -2, vjust = 2, colour = "black", size = 16, fontface = "bold")
q3 <- q4C.box + theme(legend.position = "none")  + theme(axis.title.x = element_blank())
q3 <- ggdraw(q3) + draw_plot_label(label = "C",x = 0, y = 1, hjust = -2, vjust = 2, colour = "black", size = 16, fontface = "bold")
q4 <- q4D.box + theme(legend.position = "none") + theme(axis.title.x = element_blank())
q4 <- ggdraw(q4) + draw_plot_label(label = "D",x = 0, y = 1, hjust = -2, vjust = 2, colour = "black", size = 16, fontface = "bold")

qplot <- grid.arrange(q1, q2, q3, q4, ncol = 2, nrow = 2)
qplot
ggsave(qplot, width=25, height=25, dpi = 600, units=c("cm"), filename="output/Figure4_MembranousOrganelles.pdf", bg="transparent")

```

Biplots for supplement
```{r}

# set up theme for all biplots
biplottheme <- 
  theme(panel.background = element_rect(fill = "transparent"),
        panel.grid.minor = element_blank(),
        panel.grid.major = element_blank())+
  theme(plot.background = element_rect(fill = "transparent"),
        legend.background = element_rect(fill = "transparent"),
        legend.text = element_text(colour = "black", size = 12),
        legend.title = element_text(colour = "black", size = 15),
        legend.key = element_rect(fill = "transparent")) + 
  theme(axis.text.x = element_text(face='plain', colour = "black", size = 12), 
        axis.text.y = element_text(colour = "black", size = 12),
        axis.line = element_line(colour="black", linetype = "solid"),
        axis.ticks = element_line(colour="black", linetype = "solid"),
        axis.title.x = element_text(colour = "black", size = 15),
        axis.title.y = element_text(colour = "black", size = 15)) +
  theme(legend.position = "right", legend.text = element_text(size = 12, face = "italic"))

# Supplementary Figure 1: Pseudopod relative to total cell size
# Make and save plot as pdf for plotting pseudopod area against total cell size in um^2
q1.scatter = ggplot(temdata, aes(x=sperm_surfacearea_total, y=pseudopod_surfacearea, color=species)) + 
  geom_point(alpha = 0.55, size = 3) +
  geom_smooth(method = "lm", aes(fill=species), linewidth=2) +
  labs(x = "Total sperm cell size "~(µm^2), y = "Pseudopod area "~(µm^2)) +
  scale_color_manual(values=cols2, name="species") +
  scale_fill_manual(values=cols2, name="species") +
  biplottheme 
q1.scatter 
# save as pdf
ggsave(q1.scatter, width=20, height=15, dpi = 600, units=c("cm"), filename="output/FigureS1_PseudopodPerTotalCell.pdf", bg="transparent")


# Supplementary Figure 2: Total organelle sizes
# Make and save plot as pdf for plotting total MO against total mitochondrial area in um^2
q2.scatter = ggplot(temdata, aes(x=MO_surfacearea_total, y=mitochondria_surfacearea_total, color=species)) + 
  geom_point(alpha = 0.55, size = 3) +
  geom_smooth(method = "lm", aes(fill=species), linewidth=2) +
  labs(x = "Total MO area "~(µm^2), y = "Total mitochondrial area "~(µm^2)) +
  scale_color_manual(values=cols2, name="species") +
  scale_fill_manual(values=cols2, name="species") +
  biplottheme 
q2.scatter 
# save as pdf
ggsave(q2.scatter, width=20, height=15, dpi = 600, units=c("cm"), filename="output/FigureS2_MitoAgainstMOTotals.pdf", bg="transparent")

# Assemble 2 panels
q1 <- q1.scatter + theme(legend.position = "none")
q1 <- ggdraw(q1) + draw_plot_label(label = "A",x = 0, y = 1, hjust = -2, vjust = 2, colour = "black", size = 16, fontface = "bold")
q2 <- q2.scatter + theme(legend.position = "none")
q2 <- ggdraw(q2) + draw_plot_label(label = "B",x = 0, y = 1, hjust = -2, vjust = 2, colour = "black", size = 16, fontface = "bold")

# Extract the legend from one of the original plots
legend <- get_legend(q2.scatter + theme(legend.position = "right"))

# Combine panels
supp.panels <- plot_grid(q1, q2, ncol = 2, nrow = 1, rel_widths = c(1, 1))
# Add legend
supp.panels <- plot_grid(supp.panels, legend, ncol = 2, rel_widths = c(4, 1))
# Save
ggsave(supp.panels, filename="output/SuppFigure.pdf", width=30, height=10, dpi = 600, units=c("cm"), bg="transparent")


#### Mitochondria vs. MO:

## Analysis prep
## make an indicator variable that is 0 for C.n. and 1 for C.m..
temdata$TestSpecies = as.numeric(temdata$species == "C. macrosperma")

## Analysis of difference between species for MO and mitochondrial surface area
MO_mito_model = lm(mitochondria_surfacearea_total ~ MO_surfacearea_total*TestSpecies, data = temdata)
# OR
MO_mito_model = lm(mitochondria_surfacearea_total ~ TestSpecies + MO_surfacearea_total + TestSpecies:MO_surfacearea_total, data = temdata)

summary(MO_mito_model)

## TestSpecies is significant, meaning the two species are different or rather their two regression lines differ from one another
##  MO_surfacearea_total to mitochondria_surfacearea_total above cut off, marginally significant, meaning their relationship is not significantly different from a flat line (one does not predict the other if adhering strictly to the significance cut off)
## TestSpecies:MO interaction term is not significant meaning their slopes are not significantly different from each other (if they were significant, the regression lines would be pointing in different directions)
## In cases of non-sig result with interaction term, common practice is to remove the interaction predictor and re-run the model, UNLESS we are specifically interested in the interaction. In my case, we do not care about mito vs. MO generally, BUT ONLY if they differ between species, so whether there is an interaction between species and organelle. Hence, we do not re-run the model.

## (Technically, it is saying that for C. n. but relationship in C. m is not significantly different than relationship in C. n)

## Report the following: Coefficients table as is, plus DF, plus multiple R-squared, plus adjusted R squared

# Retrieve means for total mitochondria surface area (also found in t-test results from box plots)
mean(temdata$mitochondria_surfacearea_total[temdata$species == "C. macrosperma"])
mean(temdata$mitochondria_surfacearea_total[!(temdata$species == "C. macrosperma")])

```

Other cell component comparisons
```{r Additional cell component metrics and t-test, message=FALSE, warning=FALSE}

# Total nucleus area in µm^2
q.box = ggplot(temdata, aes(species, nucleus_surfacearea, color=species)) + 
  geom_boxplot(alpha = 0, lwd = 1.3) + 
  geom_jitter(alpha = 0.55, width = 0.2, size = 4) +
  labs(x = " ", y = "Total nucleus area "~(µm^2), 
       title = "Total nucleus area") +
  scale_color_manual(values=cols2, name="species") +
  TEMtheme
q.box 

# save as pdf
ggsave(q.box, width=15, height=17.5, dpi = 600, units=c("cm"), filename="output/NucleusArea.pdf", bg="transparent")

# test differences in nucleus area (report in table: t-statistic, df, p value, confidence?, mean of x and mean of y, SE of difference)
t.test(temdata[temdata$species == "C. nouraguensis", "nucleus_surfacearea"], temdata[temdata$species == "C. macrosperma", "nucleus_surfacearea"], paired = F)

SEofCmacroNucleus<- (sd(temdata[temdata$species == "C. macrosperma", "nucleus_surfacearea"]))/sqrt(n_Cmacro)
SEofCmacroNucleus

SEofCnouraNucleus <- (sd(temdata[temdata$species == "C. nouraguensis", "nucleus_surfacearea"]))/sqrt(n_Cnoura)
SEofCnouraNucleus
#___________________________________________________

# Standardized area of mitochondria per total cell in µm^2
q.box = ggplot(temdata, aes(species, mitopertotalcell, color=species)) + 
  geom_boxplot(alpha = 0, lwd = 1.3) + 
  geom_jitter(alpha = 0.55, width = 0.2, size = 4) +
  labs(x = " ", y = "Standardized mitochondrial area per total sperm cell "~(µm^2), 
       title = "Standardized mitochondrial area per total sperm cell") +
  scale_color_manual(values=cols2, name="species") +
  TEMtheme
q.box 

# save as pdf
ggsave(q.box, width=15, height=17.5, dpi = 600, units=c("cm"), filename="output/StandardizedMitochondriaAreaPerTotalSpermCell.pdf", bg="transparent")

# test differences in standardized mitochondrial area per total sperm cell (report in table: t-statistic, df, p value, confidence?, mean of x and mean of y, SE of difference)
t.test(temdata[temdata$species == "C. nouraguensis", "mitopertotalcell"], temdata[temdata$species == "C. macrosperma", "mitopertotalcell"], paired = F)

SEofCmacroMitopertotalcell<- (sd(temdata[temdata$species == "C. macrosperma", "mitopertotalcell"]))/sqrt(n_Cmacro)
SEofCmacroMitopertotalcell

SEofCnouraMitopertotalcell <- (sd(temdata[temdata$species == "C. nouraguensis", "mitopertotalcell"]))/sqrt(n_Cnoura)
SEofCnouraMitopertotalcell
#__________________________________________________

# Density of mitochondria per cell body area
q.box = ggplot(temdata, aes(species, mitodensity, color=species)) + 
  geom_boxplot(alpha = 0, lwd = 1.3) + 
  geom_jitter(alpha = 0.55, width = 0.2, size = 4) +
  labs(x = " ", y = "Mitochondrial density per sperm cell body", 
       title = "Density of mitochondria: Number of mitochondria per area of sperm cell body "~(µm^2)) +
  scale_color_manual(values=cols2, name="species") +
  TEMtheme
q.box 

# save as pdf
ggsave(q.box, width=15, height=17.5, dpi = 600, units=c("cm"), filename="output/MitoDensity_NumberOfMitoPerAreaOfSpermCellBody.pdf", bg="transparent")

# test differences in density of mitochondria area per cell body only (no pseudopod) (report in table: t-statistic, df, p value, confidence?, mean of x and mean of y, SE of difference)
t.test(temdata[temdata$species == "C. nouraguensis", "mitodensity"], temdata[temdata$species == "C. macrosperma", "mitodensity"], paired = F)

SEofCmacroMitoDensity<- (sd(temdata[temdata$species == "C. macrosperma", "mitodensity"]))/sqrt(n_Cmacro)
SEofCmacroMitoDensity

SEofCnouraMitoDensity <- (sd(temdata[temdata$species == "C. nouraguensis", "mitodensity"]))/sqrt(n_Cnoura)
SEofCnouraMitoDensity

#___________________________________________________

# Make and save plot as png for plotting total mitochondria area against cell body (no pseudopod) in um^2
q.scatter = ggplot(temdata, aes(x=cellbodywithoutpseudopod, y=mitochondria_surfacearea_total, color=species)) + 
  geom_point(alpha = 0.55, width = 0.2, size = 4) +
  geom_smooth(method = "lm", aes(fill=species), linewidth=2) +
  labs(x = "Area of cell body w/o pseudopod [um^2]", y = "Total mitochondrial area [um^2]", 
       title = "Mitochondrial area per cell body area") +
  scale_color_manual(values=cols2, name="species") +
  scale_fill_manual(values=cols2, name="species") +
  TEMtheme
q.scatter 

# save as pdf
ggsave(q.scatter, width=25, height=17.5, dpi = 600, units=c("cm"), filename="output/MitoPerCellbody.pdf", bg="transparent")

## Analysis of difference between species for mitochondria surface area per cellbody surface area
mito_per_cell_model = lm(mitochondria_surfacearea_total ~ cellbodywithoutpseudopod*TestSpecies, data = temdata)
# OR:
#mito_per_cell_model = lm(mitochondria_surfacearea_total ~ TestSpecies + cellbodywithoutpseudopod + TestSpecies:cellbodywithoutpseudopod, data = temdata)

summary(mito_per_cell_model)

#___________________________________________________

# Standardized area of MO per total cell in µm^2
q.box = ggplot(temdata, aes(species, MOpertotalcell, color=species)) + 
  geom_boxplot(alpha = 0, lwd = 1.3) + 
  geom_jitter(alpha = 0.55, width = 0.2, size = 4) +
  labs(x = " ", y = "Standardized MO area per total sperm cell" ~(µm^2), 
       title = "Standardized MO area per total sperm cell" ~(µm^2)) +
  scale_color_manual(values=cols2, name="species") +
  TEMtheme
q.box 

# save as pdf
ggsave(q.box, width=15, height=17.5, dpi = 600, units=c("cm"), filename="output/StandardizedMOAreaPerTotalSpermCell.pdf", bg="transparent")

# test differences in standardized MO-body area per total sperm cell (report in table: t-statistic, df, p value, confidence?, mean of x and mean of y, SE of difference)
t.test(temdata[temdata$species == "C. nouraguensis", "MOpertotalcell"], temdata[temdata$species == "C. macrosperma", "MOpertotalcell"], paired = F)

SEofCmacroMOpertotalcell<- (sd(temdata[temdata$species == "C. macrosperma", "MOpertotalcell"]))/sqrt(n_Cmacro)
SEofCmacroMOpertotalcell

SEofCnouraMOpertotalcell <- (sd(temdata[temdata$species == "C. nouraguensis", "MOpertotalcell"]))/sqrt(n_Cnoura)
SEofCnouraMOpertotalcell
#___________________________________________________

# Standardized area of MO per cell body in µm^2
q.box = ggplot(temdata, aes(species, MOpercellbody, color=species)) + 
  geom_boxplot(alpha = 0, lwd = 1.3) + 
  geom_jitter(alpha = 0.55, width = 0.2, size = 4) +
  labs(x = " ", y = "Standardized MO area per sperm cell body" ~(µm^2), 
       title = "Standardized MO area per sperm cell body" ~(µm^2)) +
  scale_color_manual(values=cols2, name="species") +
  TEMtheme
q.box 

# save as pdf
ggsave(q.box, width=15, height=17.5, dpi = 600, units=c("cm"), filename="output/StandardizedMOAreaPerCellBody.pdf", bg="transparent")

# test differences in standardized MO-body area per cell body only (no pseudopod) (report in table: t-statistic, df, p value, confidence?, mean of x and mean of y, SE of difference)
t.test(temdata[temdata$species == "C. nouraguensis", "MOpercellbody"], temdata[temdata$species == "C. macrosperma", "MOpercellbody"], paired = F)

SEofCmacroMOpercellbody<- (sd(temdata[temdata$species == "C. macrosperma", "MOpercellbody"]))/sqrt(n_Cmacro)
SEofCmacroMOpercellbody

SEofCnouraMOpercellbody <- (sd(temdata[temdata$species == "C. nouraguensis", "MOpercellbody"]))/sqrt(n_Cnoura)
SEofCnouraMOpercellbody
#___________________________________________________

# Number of MOs per idealized cell body circumference
q.box = ggplot(temdata, aes(species, MOpercircumference, color=species)) + 
  geom_boxplot(alpha = 0, lwd = 1.3) + 
  geom_jitter(alpha = 0.55, width = 0.2, size = 4) +
  labs(x = " ", y = "Number of MOs per idealized sperm cell circumference", 
       title = "Number of MOs per idealized sperm cell circumference") +
  scale_color_manual(values=cols2, name="species") +
  TEMtheme
q.box 

# save as pdf
ggsave(q.box, width=15, height=17.5, dpi = 600, units=c("cm"), filename="output/NumberMOsPerCellBodyCircumference.pdf", bg="transparent")

# test differences in number of MOs area per idealized cell body circumference (report in table: t-statistic, df, p value, confidence?, mean of x and mean of y, SE of difference)
t.test(temdata[temdata$species == "C. nouraguensis", "MOpercircumference"], temdata[temdata$species == "C. macrosperma", "MOpercircumference"], paired = F)

SEofCmacroMOpercircumference<- (sd(temdata[temdata$species == "C. macrosperma", "MOpercircumference"]))/sqrt(n_Cmacro)
SEofCmacroMOpercircumference

SEofCnouraMOpercircumference <- (sd(temdata[temdata$species == "C. nouraguensis", "MOpercircumference"]))/sqrt(n_Cnoura)
SEofCnouraMOpercircumference

#___________________________________________________

# Make and save plot as png for plotting total MO area against cell body (no pseudopod) in um^2
q.scatter = ggplot(temdata, aes(x=cellbodywithoutpseudopod, y=MO_surfacearea_total, color=species)) + 
  geom_point(alpha = 0.55, width = 0.2, size = 4) +
  geom_smooth(method = "lm", aes(fill=species), linewidth=2) +
  labs(x = "Area of cell body w/o pseudopod [um^2]", y = "Total MO area [um^2]", 
       title = "MO area per cell body area") +
  scale_color_manual(values=cols2, name="species") +
  scale_fill_manual(values=cols2, name="species") +
  TEMtheme
q.scatter 

# save as pdf
ggsave(q.scatter, width=25, height=17.5, dpi = 600, units=c("cm"), filename="Figure2A_MOPerCellbody.pdf", bg="transparent")

## Analysis of difference between species for MO surface area per cellbody surface area
MO_per_cell_model = lm(MO_surfacearea_total ~ cellbodywithoutpseudopod*TestSpecies, data = temdata)

summary(MO_per_cell_model)

#___________________________________________________

# Make and save plot as pdf for plotting number of MOs per idealized cell circumference
q.scatter = ggplot(temdata, aes(x=IdealizedCircumference, y=MO_count, color=species)) + 
  geom_point(alpha = 0.55, width = 0.2, size = 4) +
  geom_smooth(method = "lm", aes(fill=species), linewidth=2) +
  labs(x = "Idealized circumference of cell body without pseudopod", y = "MO count", 
       title = "MO count per idealized cell circumference w/o pseudopod") +
  scale_color_manual(values=cols2, name="species") +
  scale_fill_manual(values=cols2, name="species") +
  TEMtheme
q.scatter 

# save as pdf
ggsave(q.scatter, width=25, height=17.5, dpi = 600, units=c("cm"), filename="Figure2A_MOCountPerCircumference.pdf", bg="transparent")

## Analysis of difference between species for MO-body number per idealized cell body circumference
MO_per_circumference_model = lm(MO_count ~ IdealizedCircumference*TestSpecies, data = temdata)

summary(MO_per_circumference_model)

```
````
